# Precise age estimation in clonal species using a somatic genetic clock

**DOI:** 10.1101/2023.11.07.566010

**Authors:** Lei Yu, Jessie Renton, Agata Burian, Marina Khachaturyan, Jonne Kotta, John J. Stachowicz, Katherine DuBois, Iliana B. Baums, Benjamin Werner, Thorsten B. H. Reusch

## Abstract

Age and longevity are key parameters for demography and life-history evolution of organisms. In clonal species, a widespread life history among animals, plants, algae and fungi, the sexually produced offspring (the genet) grows indeterminately by producing iterative modules, or ramets. The age of large genets often remains elusive, while estimates based on their spatial extent as proxy for age are unreliable. Here, we present a method for age estimation using a molecular clock based on the accumulation of fixed somatic genetic variation (SoGV) that segregates among ramets of the same genet. Using a stochastic model of a generic clonal organism, we demonstrate that the accumulation of fixed SoGV via somatic genetic drift will approach linearity after a short lag phase, and is determined by the mitotic mutation rate, without direct dependence on asexual generation time. The lag phase decreased with lower stem cell population size (*N*), number of founder cells for the formation of new modules (*N*_*0*_), and the ratio of symmetric vs. asymmetric stem cell divisions. We apply the somatic genetic clock to the clonal plant model *Zostera marina* (eelgrass) and show that linearity is approached within a few years. Taking advantage of two long-term cultivation experiments for *Z. marina* (4 and 17 years respectively) as calibration points, we find genet ages up to 1,403 years in a global data set of 20 eelgrass populations. The somatic genetic clock is applicable to any multicellular clonal species where a small number of founder cells are recruited to form new ramets, opening novel research avenues to study longevity and hence, demography and population dynamics of clonal species.

## Introduction

Clonal reproduction is the process of generating (potentially) physically independent multi-cellular organisms (i.e., ramets *sensu* ref^1^ via mitosis, a widespread life-history among animals, plants, algae and fungi^2^. Starting from a single zygote, multipotent somatic cells proliferate to form new ramets via branching or budding, often becoming physiologically independent after a few years when severing from the parental tissue. All modules or ramets stemming from that single zygote represent a genet (or clone). Often, the contribution of sexual and clonal reproduction to local population structure varies among species and localities^3-5^, resulting in asexual populations of ramets that are nested within the “classical” population of genets^2,6^. Coral, algae, seagrass, or poplar genets, for example, can reach considerable size and therefore age with linear extents of >1 km^7-11^. The apparent persistence and resilience of asexual ramet populations is astonishing in light of the considerable temporal and spatial variation they may experience over their lifetimes despite little genetic variation (but see refs^10,12^) and raises questions about these species’ adaptability in a rapidly changing climate^13^.

As a key parameter to evaluate this persistence, genet age/longevity has been inherently difficult to estimate, in particular, when biomass tracing back to an individual’s origin is not preserved, as is the case in non-woody plants^14^. For example, a small genet is not necessarily young if episodes of ramet mortality reduced its size in the past. In order to estimate genet age via molecular genetic methods, somatic genetic variation segregating among ramets has previously been used. However, those attempts lacked resolution, as the somatic genetic variation could be estimated at only a few marker loci^9,15^.

Here, we present a novel approach to estimate genet age based on a somatic genetic clock that uses complete genome information of the focal species. Molecular clocks were initially developed for species-level phylogenies and rely on the neutral theory of molecular evolution^16^. Fixed neutral mutations within species accumulate at a constant rate equal to the rate of spontaneous mutations per unit time Easteal, 1988 #4789}, and thus genetic differences between species increase with absolute time^17,18^. If the mutation rate can be derived based on calibration points such as fossil evidence, clock estimates can be extended to phylogenetically related clades^19^. Recently, fixation of somatic genetic variation (hereafter SoGV) was demonstrated in clonal species through a process of somatic genetic drift^12^. During genet growth via new ramet formation, somatic mutations become fixed in the descendant ramets, essentially because only a few pluripotent cells of the proliferating tissue are recruited to form the new module or ramet^12,20,21^ (Fig. 1, Supplementary Fig. 1). Here, we built upon these findings and introduce the somatic genetic clock that uses the rate of genome-wide, asexual fixation of alleles to estimate the extent of differentiation between the founder and descendant ramets of a genet. In doing so, we can infer the time to the least common ancestor of multiple or pairs of ramets, here the zygote, and derive a “somatic genetic clock” that permits the precise ageing of large plant clones (genets) and possibly, other clonal animal, algal or fungal species.

**Figure 1:**
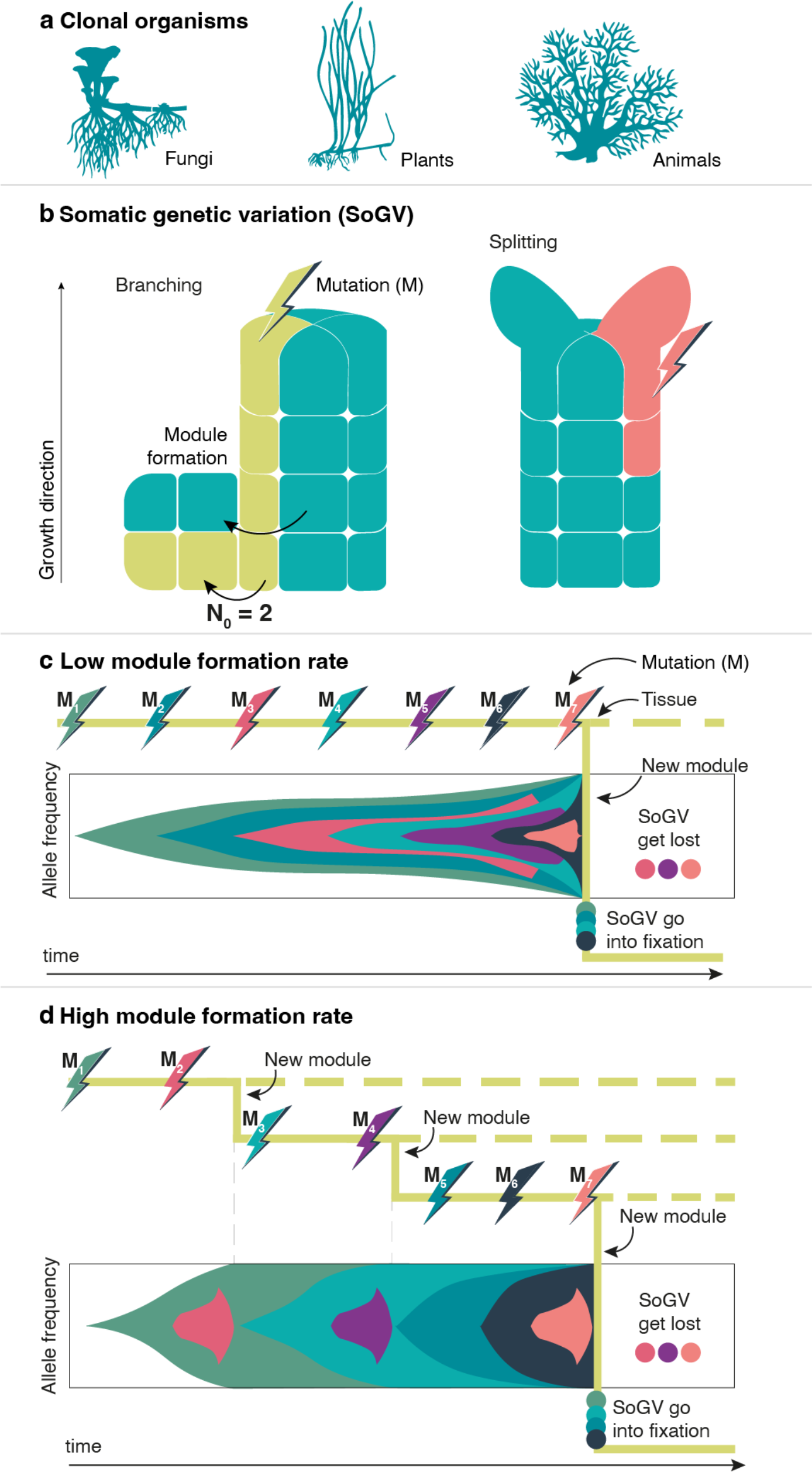
Dynamics of somatic genetic variation (SoGV) in generic clonal organisms. **a**, Multicellular clonal species exist across the tree of life. **b**, Allele frequency change of SoGV due to the formation of new modules by branching or splitting. A new module is initiated either directly by the stem cells (i.e., splitting), or by the daughter cells of the stem cells (i.e., branching). Splitting reduces the size of the original stem cell population, while branching leaves the original cell population untouched. During the formation of new modules, the cell population undergoes a genetic bottleneck. **c, d**, The accumulation rate of fixed SoGV is independent of module formation rate. The tree topology depicts a module undergoing (multiple) module formation events, after which the dashed line and the solid line represent the original module and the new module respectively. New mutations (M) occur at a constant rate, and only mutations in the new modules are depicted (with a different color). For each time point, the vertical length of the colors represents the frequency of the SoGV within the module. Clonal dynamics in a single module (solid line in tree structure) are depicted as a Muller plot which shows the nested allele frequency of SoGV over time. The frequency of SoGV changes during module formation events, due to the bottleneck. Eventually, SoGV are either fixed or lost. For **c**, low module formation rate, fixation events are rare. Thus, many SoGV have accumulated in the intervening time and are fixed simultaneously. In **d**, under high module formation rate, fixation events occur more frequently, but with fewer SoGV fixed at each branching event.

## Results

### A generic somatic genetic clock in clonal species revealed by modelling and simulations

To estimate the time over which fixed somatic genetic variation (SoGV) accumulates and segregates under clonal growth, we developed a stochastic, agent-based model of a generic clonal organism that comprises a collection of modules, adapted from population genetics models of cancer evolution^22^ (Material & Methods). Within this model, a module is simplified to the stem cell population of a single ramet (all somatic cells are derived from stem cells, and thus, can be ignored). Cells and modules are subject to stochastic update events including cell division, death, and the formation of new modules, with new Poisson-distributed mutations occurring at each cell division. We considered a range of scenarios with different types of stem cell division (symmetric vs. asymmetric), different (founder) stem cell pool sizes, and varying rates and mechanisms for forming new modules (branching vs. splitting), attempting to capture possible life history variation in clonal species across the tree of life (Fig. 2). We found that, given sufficient time, any scenario would converge to a constant accumulation rate of fixed SoGV, and thus the number of fixed SoGV would increase linearly with clonal age (Fig. 3, Supplementary Figs. 2,3) as required for a useful molecular clock.

**Figure 2:**
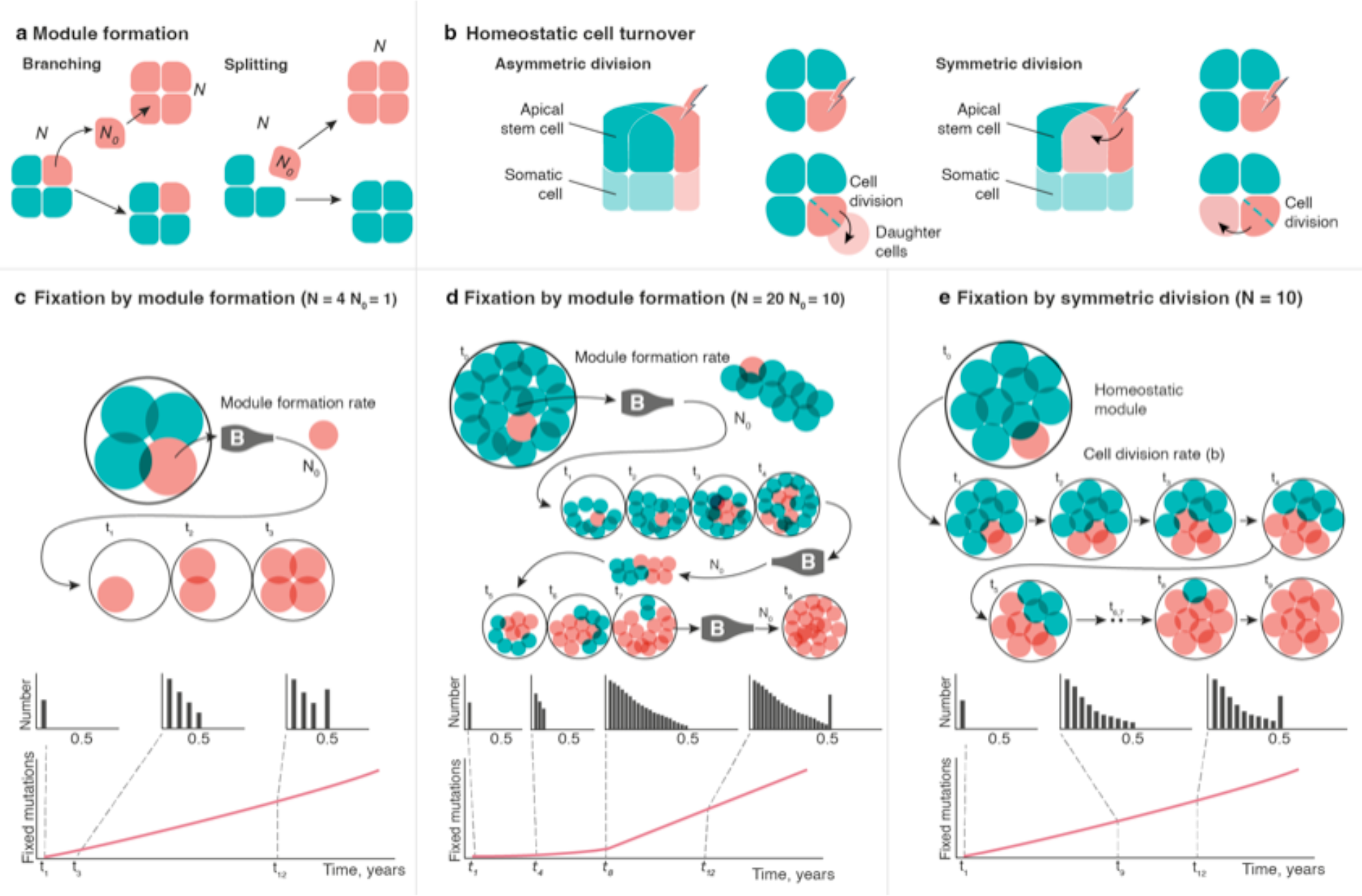
Processes determining fixation rates of somatic genetic variation in a generic clonal organism. Modules represent the stem cell compartment of each ramet within the clonal organism. **a**, New module formation occurs by branching or splitting of homeostatic modules, during which *N*_*0*_ cells are selected from a parent module to form the founding population of a new module. For module branching these are copied (leaving the parent unchanged), while for module splitting, they are removed from the parent. Growth by cell division is then implemented so that all modules return to homeostatic size *N*. **b**, Cell turnover in homeostatic modules occurs by asymmetric or symmetric division of stem cells. Dashed lines depict dividing stem cells that produce two daughter cells. After asymmetric division, one daughter cell remains in the module and the other differentiates (leaving the stem cell compartment). Under symmetric division, one daughter cell replaces one of the other stem cells (which is assumed to differentiate), and thus both daughter cells remain in the module. **c-e**, The frequency of a new mutation within the stem cell population (*N*) is 1/(2\**N*) for diploid species. This frequency will change during clonal proliferation. If not lost by drift, persistent mutations can be visualized based on their frequencies relative to the total number of chromosomes in the stem cell population, i.e., 1/(2\**N*), 2/(2\**N*), …, *N*/(2\**N*). A frequency of *N*/(2\**N*)=0.5 means the mutation is shared by all the stem cells, reaching the fixation state (i.e., fixed SoGV). The number of fixed SoGV accumulates linearly in modules once an equilibrium is reached. **c, d**, SoGV become fixed in modules by repeated bottlenecks (depicted as a bottle labelled “B”) induced by module formation (i.e. module size reduces from *N* to *N*_*0*_ then regrows). The time period of the non-linear phase is shorter for **c**, smaller *N* and *N*_*0*_ compared to **d**, larger *N* and *N*_*0*_. **E**, SoGV become fixed in modules during homeostasis in the case of symmetric division. This is similar to classic population genetic models (i.e. a Moran process), and the time period of the non-linear phase increases with *N*.

**Figure 3:**
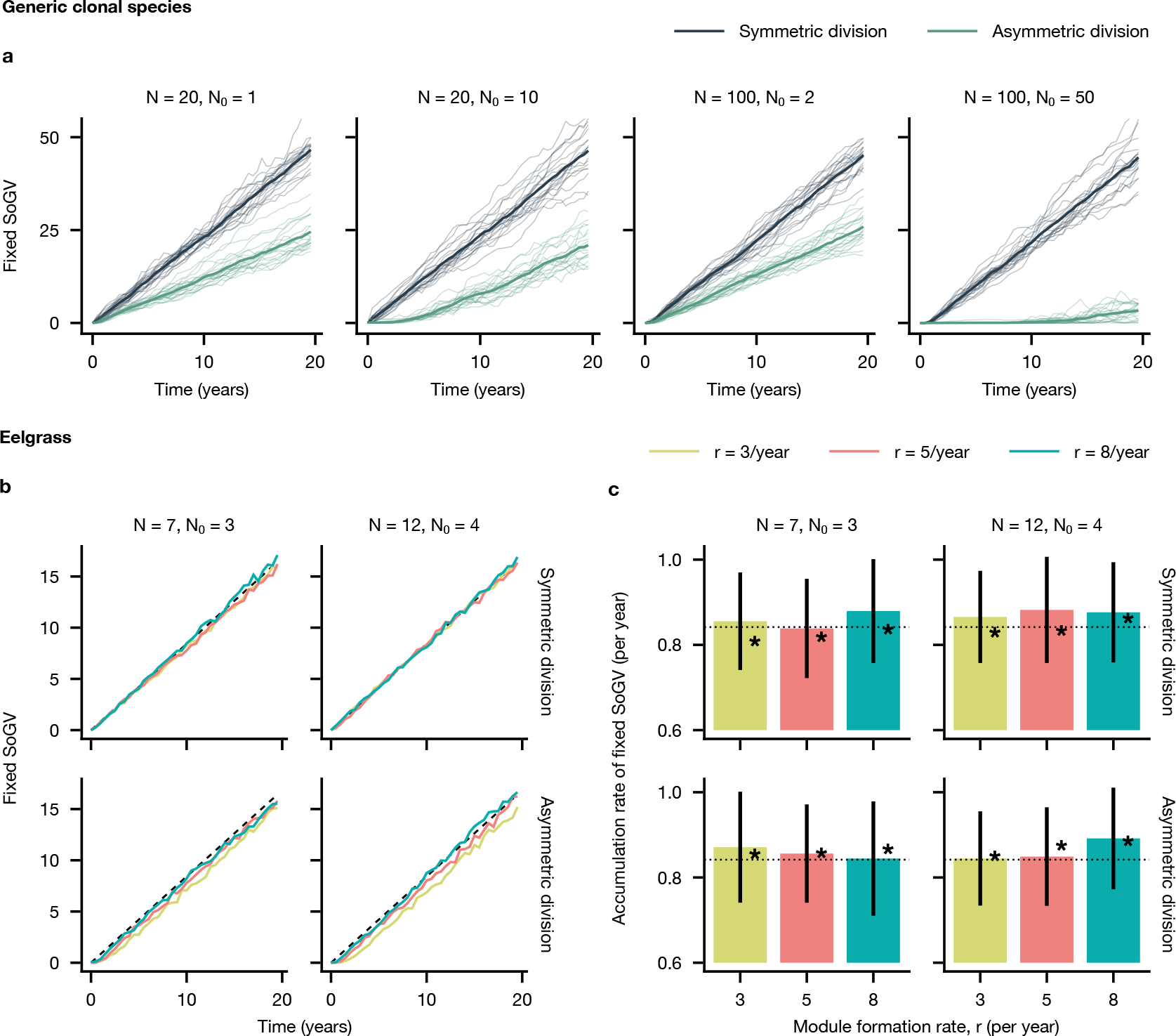
Agent-based model predictions for the accumulation of fixed somatic mutations via somatic genetic drift. **a**, Model for a generic clonal species. Simulations are shown for a range of model regimes, with new modules formed by module branching. After a lag phase the rate of accumulation of fixed SoGV reaches an equilibrium and becomes linear. However, in some cases (N=100; N_0_=50, asymmetric division), the lag phase is long. Thin lines: mean over all modules of a single genet; thick line: mean over all twenty genets. Chosen parameters: μ=0.01, *b*=122/year, *r*=5/year, Z=100 (symmetric and asymmetric update events occur at rate b). **b, c**, Model parametrization for eelgrass (Supplementary Figures 7-9), demonstrating that the equilibrium rate of accumulation of fixed SoGV is reached quickly. New modules are formed by module branching, which is closer to the biological process in eelgrass. **b**, Data are means over ten simulations. In each simulation the mean fixed mutations are calculated at each time point from a random sample of ten modules. Dashed line: μb (approximation of the mutation rate per cell per year). **c**, The accumulation rate of fixed SoGV is estimated from simulated modules at two time points mimicking the experimental methodology (4 years: 3 clones with 2 sampled modules per clone, and 17 years: 2 clones with 5 and 6 sampled modules respectively; see Materials & Methods). Mean fixed SoGV is calculated for each clone and the accumulation rate is then estimated by linear regression. Bars and error bars: mean and standard deviation, respectively, from 100 repeats; dashed line: μb; stars: accumulation rate of fixed SoGV is estimated by performing linear regression on 100 simulated 200-year old clones and taking the mean. Parameters: μ=0.0069, *b*=122/yr, *Z*=1000 (symmetric update events occur at a rate *b*/2, asymmetric update events at rate *b*).

The accumulation rate of fixed SoGV was solely determined by the mutation rate per cell per site per year. While the module formation rate (*r*) does not directly impact the accumulation rate of fixed SoGV (Fig. 1), it can have a small indirect effect by altering the mutation rate, either as a result of stochasticity or because of different effective mutation rates during homeostasis and growth. This effect is small (Supplementary Fig. 4) and we consider it negligible for biologically relevant parameter ranges. The relative constancy despite different module formation rates, i.e. asexual generation times, is equivalent to the classical molecular clock being dependent only on mutation rate and not sexual generation time^18,23,24^.

**Figure 4:**
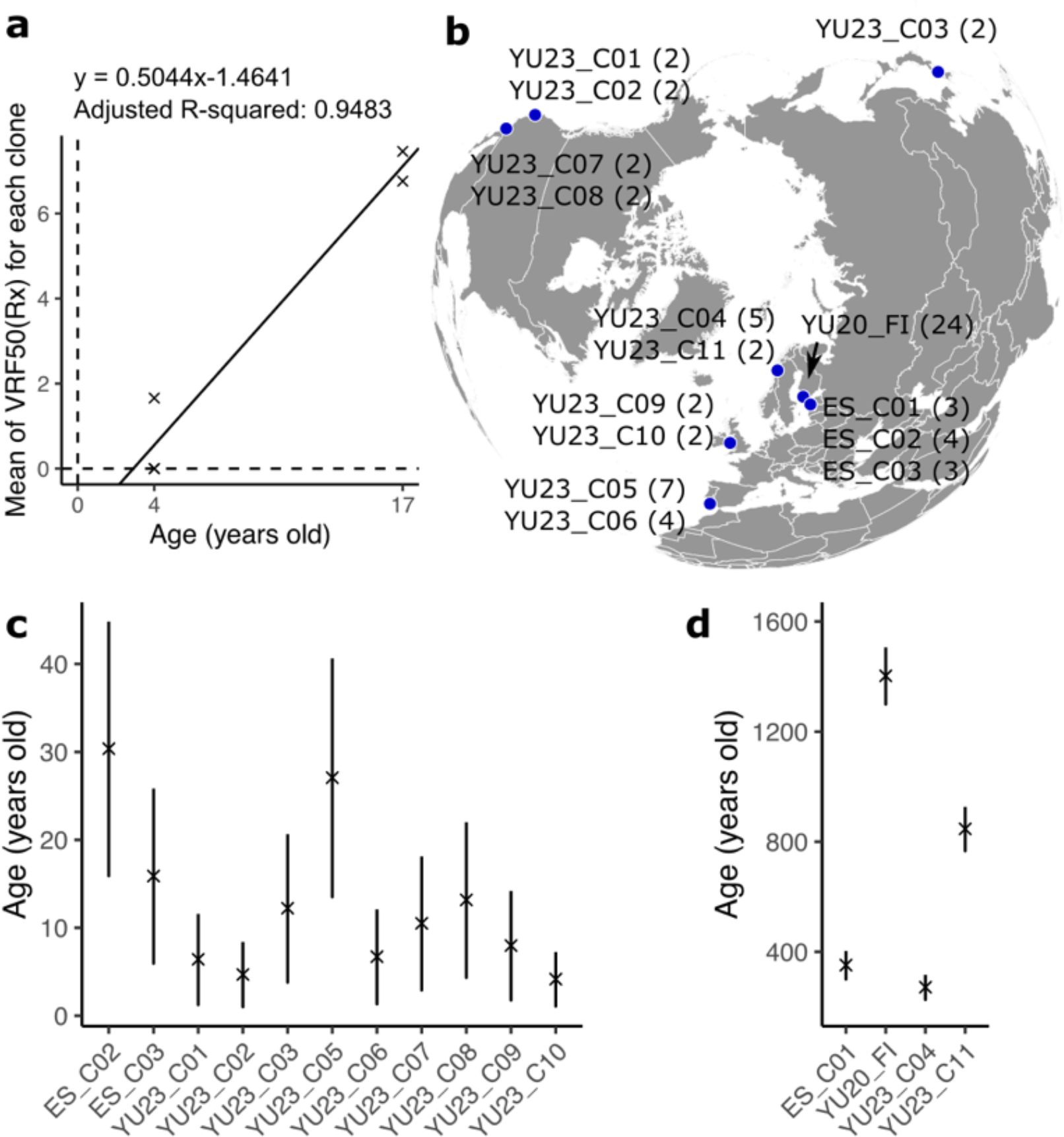
Estimating the age of globally distributed eelgrass (*Zostera marina*) clonal lineages based on the somatic genetic clock. **a**, Accumulation rate of the VRF50(R_x_). Each data point represents one clonal lineage. Two of the three 4-yr-old clonal lineages show mean VRF50(R_x_) of 0, and the data points overlap with each other. **b**, Location of the 15 globally distributed eelgrass genets. Number in parentheses indicates the number of ramets for each corresponding genet. **c**, Age estimates for relatively young genet. Data point indicate the age estimate with the 95% confidence interval. **d**, Age estimates for the 4 oldest genets. The detailed information for age estimates available in Supplementary Data 1.

We next explored the duration of the lag-phase before linearity is reached and found that it depended upon the size of the stem cell pool per module (*N*), the number of founder stem cells that are recruited to form new modules (*N*_*0*_), the ratio of symmetric *vs*. asymmetric cell division, the rate of stem cell division (*b*), the rate at which new modules are formed (*r*) and whether they are formed by branching or splitting (Fig. 3a, Supplementary Figs. 5,6). Module formation via a small number of founder stem cells (small *N*_*0*_) reached a linear equilibrium fast for both branching and splitting (Fig. 3a, Supplementary Fig. 6). The duration of the lag-phase increased substantially for a large number of founder cells and/or solely asymmetric stem cell divisions. Fixation of SoGV occurs due to the repeated formation of new modules, during which the population of cells that form the module undergo a bottleneck (Fig. 2). Additionally, fixation can occur due to homeostatic cell turnover within the module if, and only if, there is symmetric cell division while this cannot occur for purely asymmetric divisions. Next, we estimated the conditional fixation times for different clonal species’ life-histories. Assuming asymmetric cell division, fixation occurs only due to repeated module formation, which can be represented as a modified Wright-Fisher process. We derive the conditional fixation times, which are approximately 4*N*_0_(1 − *N*_0_/*N*)/*r* (Eq. 1) for module splitting and 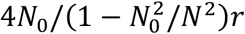 (Eq. 2) for module branching (see Supplementary Note 1.3 for the derivation using a diffusion approximation). Thus, fixation times may be decreased by reducing *N*_*0*_, even when *N* is large (Supplementary Fig. 5). For symmetric cell division, fixation due to homeostatic cell turnover usually dominates, because the cell division rate *b* is greater than the module formation rate *r*. The conditional fixation time is therefore better represented by a Moran process, approximately *N/b* (Eq. 3) (ref^25^). The conditional fixation time can be considered as a lower-bound on the lag-phase to reach equilibrium accumulation of fixed SoGV. Thus, these equations indicate the absolute timescale over which the somatic genetic clock is applicable for different species life-histories.

### Application of the somatic genetic clock in eelgrass *Z. marina*

We then applied the somatic genetic clock to the seagrass *Zostera marina* (eelgrass), an emerging model for evolution in clonal plants. We first examined the structure of the shoot apical meristem (SAM) containing a population of stem cells in higher plants^26^ via laser confocal microscopy. We were interested in evidence for SAM stratification, the likely number of stem cells (*N*) and module founder cells (*N*_0_), as well as the stem cell division mode (symmetric or asymmetric) (Supplementary Note 2). We found that the SAM was organized into one-layered L1 (tunica) and underlying L2 (corpus) as in many other monocotyledonous plant species (Supplementary Fig. 7a). No periclinal cell division in L1 was observed during the formation of axillary meristems, indicating a stable boundary between L1 and L2 (Supplementary Fig. 7b-d). In contrast, frequent periclinal cell divisions in L1 were observed during the formation of leaves, which suggested that L1 mostly or exclusively contributed to leaves (Supplementary Fig. 8). A likely number of L1 stem cells is between 7 and 12 with possible both asymmetric and symmetric cell division modes (Supplementary Fig. 9). From this population, about 3 or 4 stem cells give rise to cells which form a new module.

Next, we addressed how a SoGV can become fixed throughout the entire tissue of a new module despite meristem stratification. Indeed, we find clear allele fixation at *f*=0.5 in variant frequency diagrams (e.g., >7000 with *f*=0.5; ref^12^, and Supplementary Fig. 10). Although shoot meristems are generally stratified in *Z. marina* as in other angiosperms^27^ (Supplementary Fig. 7), it cannot be excluded that infrequent periclinal cell divisions occur in the L1(ref^28^) leading to SoGV fixation in all tissues. Note that leaf tissues that are derived exclusively from L1 (Supplementary Fig. 8) were predominating in the sample used for bulk sequencing. We thus continued by simplifying the fixation dynamics by assuming a one-layer case, enabling the application of our model of a generic clonal organism to eelgrass.

We parametrized the model for eelgrass and focused on the most likely range with *N*=7-12 and *N*_0_=3-4 (Fig. 3), but also considered more extreme scenarios ranging from the strongest (*N*=7, *N*_0_=1) to the weakest (*N*=12, *N*_0_=6) intensity of somatic genetic drift, in combination with branching rates 3-8/yr (ref ^29,30^). The accumulation rate of fixed SoGV remained similar (Fig. 3b & Supplementary Figs. 11a,12), indicating that mutation accumulation on the size of the shoot apical meristem (SAM) and rate of asexual reproduction was negligible.

Using Eqs. (2) and (3) we estimated the conditional fixation times for novel mutations under asymmetric and symmetric cell division, respectively, within an eelgrass clone. For the most likely parameter range these gave reasonable lower and upper bounds of 2 years (*N*=7, *N*_0_=3, *r*=8/yr) and 6 years (*N*=12, *N*_0_=4, *r*=3/yr) for asymmetric cell division, and 0.05 years (*N*=7, *b*=122/yr) and 0.1 years (*N*=12, *b*=122/yr) for symmetric division. This suggests that a constant accumulation rate required for the somatic genetic clock will be reached relatively fast in eelgrass, in the order of years or even months. This is verified by our simulations (Fig. 3b) in which we observe very small lag-times (≲1 year) for symmetric cell division. For asymmetric cell division it took longer to reach an equilibrium, with the time increasing for smaller module formation rate (*r*) and larger (founder) module size. However, the lag-times still appeared in the order of years, rather than decades.

### Calibration of the somatic genetic clock using experimental data

Next, two long-term cultivation experiments with *Z. marina* genets of known age (4 and 17 years, respectively) allowed for a calibration of the somatic genetic clock. Owing to statistical noise in estimating the true allele frequency via mapped reads at a given locus, differentiating between mosaic and fixed SoGV is inherently difficult. Hence, we developed the variable “Variant Read Frequency 50 (R_x_)” (hereafter VRF50(R_x_)) as a proxy for the number of fixed SoGV in ramet “R_x_” relative to the founder of the genet (Materials and Methods, Supplementary Fig. 13). The mean VRF50(R_x_) of a ramet population can be used as estimator for its genet age. In order to calibrate the somatic genetic clock for *Z. marina*, genets of known ages (4 and 17 yrs) were deep-sequenced (~1000x, and ~80x for 4 and 17 years, respectively) to calculate the accumulation rate of VRF50(R_x_). The mean VRF50(R_x_) and the age of a genet were used to fit a linear model (Fig. 4a, y = 0.5044x-1.4641, adjusted *R*^2^: 0.9483, *P*<0.001). To verify that our data could be used to accurately calibrate the clock, we recreated the sampling strategy for both time points, i.e. 4 and 17 yrs, by simulation and estimated the accumulation rate of fixed SoGV. All parameter settings led to similar estimated rates, which were close to the true rate (Fig. 3c). Even in the most extreme case (*N*=12, *N*_0_=6, *r*=3/yr) the accumulation rate will only be slightly underestimated (approximately 8% comparing with the estimated rate from 200-year simulations, i.e. stars in Fig. 3c and Supplementary Fig. 11b). Thus, we considered that our data could be safely used for calibration.

### Age estimation of 15 globally distributed *Z. marina* genets

We then used the calibrated somatic genetic clock to estimate the age of eelgrass genets in a worldwide data set^31^ (Fig. 4b, Supplementary Data 1). Among the 15 genets with 2 or more ramets sampled, most were <40 years old (Fig. 4c), while four attained >270 yrs (Fig. 4d), one in Estonia (352 yrs), two in Norway (271 and 847 yrs), and one in Finland (1,403 yrs). All genets >270 yrs of age were located in higher latitudes (>50°N) in the North Atlantic, indicating that marginal populations were more likely to maintain old genets^4^, and supporting the long-standing geographic parthenogenesis hypothesis^32^. Although the evolutionary history in the Pacific is much longer than that in the Atlantic^31^, Pacific eelgrass genets were young (<40 yrs). In addition, the old clonal lineages were distributed in the locations that were recolonized by glacial refugia after the last glacial maximum, indicating that clonal reproduction is a particularly successful reproductive mode to rapidly colonize newly opened areas^4^. Note that age estimates based on spatial extent would have been misleading, as genets with small spatial extent were found to be >300 yrs old. For example, while clone ES_C01 in Estonia only contained 3 ramets spreading ~20 m (Supplementary Fig. 14), it was estimated to be 352 yrs old based on the somatic genetic clock.

## Discussion

We present a somatic genetic clock that permits the precise age estimates of genets in clonally growing plants, and possibly, many clonal animal, fungal and algal species. The duration of the lag time before the DNA-sequence based somatic genetic clock approaches linearity decreases for fewer stem cells and founder stem cells; for symmetric, rather than asymmetric cell divisions; and for increased rates of new module formation. Hence, an application of the somatic genetic clock is most accurate for estimating clonal age if the stem cell population size *N* is small and new module formation happens through a small founder cell population *N*_0_ as realized in plant shoot apical meristems. In organisms that asexually reproduce through budding, time to linearity will depend on the number of cells contributing to the new bud. Conversely, marine invertebrates or algae that propagate asexually through fission will have an exceedingly long lag-time, as essentially half of all body cells comprise the founder cell population *N*_0_. By applying our analytical results (Eqs. 1-3), we are able to estimate the timescale over which the somatic genetic clock is applicable for any given organism.

Once linearity is reached, the rate of the somatic genetic clock is constant across module formation rates, thus asexual generation times, which is the hallmark of a valid molecular clock. Similar to the rate constancy despite different generation times in species-level phylogenies^23,24^, under a higher module formation rate, fewer mutations are fixed by any single module formation event, but the total number of module formation events is higher (Fig. 1), and vice versa. Our proposed clock is analogous to mitotic evolution in non-modular species, such as humans, specifically the emergence of genetic heterogeneity among healthy and cancerous human somatic tissues within an individual^33,34^. Fixed mutations within specific human tissues accumulate linearly with age^35,36^; similarly, we find that the number of fixed somatic genetic variation (SoGV) between founder and descendant ramets also accumulates at a constant rate.

Our findings on fixation processes will also apply to an evolutionary epigenetic clock that was recently described for self-fertilizing and clonally reproducing plants^37^. This clock uses the much faster accumulation of neutral gene body (de)methylations of cytosine nucleotides. As an additional step, the identification of genomic regions with clock-like behavior of (de)methylation is required^37^. The somatic genetic clock proposed here is complementary and will be best suited for slightly longer time intervals of >10 years to potentially tens of thousands of years, and where methylation data are unavailable. Here, we provide the theoretical foundation of stem cell population genetics, why both, the somatic genetic clock, and the evolutionary epigenetic clock^37^ are ultimately determined by mutation rate, as is the case for general molecular clocks^23^.

Some of the analogies of our modelled and observed temporal dynamics with classic population genetics are instructive. In our study, the stem cell population size, and the time period between two adjacent branching events, correspond to the population size *N*e, and generation time in classic population genetics, respectively. Due to the usually large *N*e (>100) in combination with genetic exchange among lineages, classic molecular clocks are limited to macro-evolutionary timescales (~10^5^-10^8^ years). However, the stem cell population size in plants is extremely small (e.g., 7-12 for eelgrass, but for other angiosperms often only 3-4, ref^26^), and module formation events often occur multiple times per year, which makes somatic genetic clock solid for recent time scales. Note that the time until stem cell populations are “saturated” with standing genetic variation, resulting from novel mutations, increases with population size *N*e, similar to time-lags required for a population to reach mutation-drift equilibrium in population genetics^38^.

With increasing availability of full genome data at the population level, our study provides an achievable and accurate method for estimating the age of clonal plants, and possibly, other clonal species in the animal and fungal kingdom^2^. It opens multiple new research avenues to model the demography, resilience and evolution of the many species that are facultatively clonal, and where direct and precise ageing information was previously unavailable.

## Methods

### Simulating fixed mutation accumulation in a clonal organism

We implemented a stochastic, agent-based model of a clonal organism, adapted from population genetics models of cancer evolution^22^. The organism is represented as a population of modules that grows to a fixed size *Z* by producing new modules via module splitting or branching. Modules consist of stem cells and have different dynamics depending on whether they are in growth or homeostasis. During the growth phase the module grows by cell division, which is implemented by a stochastic pure-birth process with rate *b*. Once the module reaches size *N* it enters homeostasis. Cell divisions are coupled with cell deaths, so that the population size remains constant. This is done either by implementing an asymmetric update (a cell divides producing only one progeny) or a symmetric update (a cell divides producing two progeny and another cell is removed from the module). This symmetric update corresponds to a Moran process. Dividing cells acquire novel, Poisson-distributed mutations with mean *μ*.

Homeostatic modules produce new modules at rate *r*. This is done by module splitting or module branching. For module splitting, the parent module donates *N*_*0*_ cells to the new child module. Both parent and child modules then re-enter the growth phase. For module branching, *N*_*0*_ cells are sampled without replacement from the parent module and then copied to form the child module which enters a growth phase. The parent module is unchanged. If the population of modules has reached maximum size *Z*, a randomly selected module is killed whenever a new module is formed to keep the population size constant.

Code is available at https://github.com/jessierenton/SomaticEvolution.jl. The simulation is implemented using a Gillespie algorithm^39^:

1. Initialize the simulation with one module that is formed of a single cell, t=0.
2. Calculate the transition rates for all transitions:
  a. Cell division in a growing module: r_a_ = bn_growth_
  b. Symmetric division in a homeostatic module: r_b_ = λNZ_homeostatic_
  c. Asymmetric division in a homeostatic module: r_c_ = γNZ_homeostatic_
  d. New module formation: r_d_ = rZ_homeostatic_
  Here, *n*_growth_ is the total number of cells in growing modules and *Z*_homeostatic_ is the number of homeostatic modules. We set λ = b (or b/2), γ = 0 for purely symmetric division and λ = 0, γ = *b* for purely asymmetric division.
3. Transition *i* is chosen with probability r_*i*_/(r_a_ + r_b_ + r_c_ + r_d_). If a cell division occurs during any transition, the newly divided cells acquire *M* ∼ Poisson(µ) novel mutations. Possible transitions are:
  a. Choose a cell to divide uniformly at random from all cells in growing modules.
  b. Choose a homeostatic module, uniformly at random. From that module choose a cell to divide and a different cell to remove, uniformly at random (Moran update).
  c. Choose a homeostatic module, uniformly at random. From that module select a cell to divide, also uniformly at random. One progeny cell remains in the module the other is removed (asymmetric division).
  d. Choose a homeostatic module uniformly at random to be the parent module and if *Z* = *Z*_max_, choose a second module to die. A new module is formed from the parent module by (i) splitting or (ii) branching. First, select *N*_*0*_ cells from parent module without replacement, then,
    i. Module branching: copy them to form a new module, leaving the parent module unchanged.
    ii. Module splitting: remove them from the parent module to form a new module.
4. Update the time *t*’ = *t* + *δt*, where *δt*~ExponentialL1/(*r*_*a*_ + *r*_*b*_ + *r*_*c*_ + *r*_*d*_)).
5. Repeat steps 2-4 until *t* = ***T***_max_.

Data is generated at discrete time-steps for the number of fixed SoGV in each module.

**Table 1.** Model parameters.

| Symbol | Description |
| --- | --- |
| $b$ | Cell division rate during growth |
| $\lambda$ | Symmetric division (Moran) rate during homeostasis |
| $\gamma$ | Asymmetric division rate during homeostasis |
| $r$ | Module formation rate |
| $N$ | Homeostatic module size (number of stem cells) |
| $N_0$ | Initial module size (number of founder stem cells) |
| $Z$ | Population size (maximum number of modules) |
| $\mu$ | Mutation rate per cell per division |

### Shoot Apex Preparation and Imaging in Laser Confocal Microscope

*Zostera marina* plants collected in Falckenstein, Kiel Fjord (54.392N, 10.192E) were kept at 8-12°C temperature and 150 µmol quanta*s^-1^*m^-2^ light intensity in 800-L indoor wave tanks, the “Zosteratron”, receiving ambient Baltic seawater while rooted in ambient sediment (12 cm deep), with an intake pipe 10km distant from the collection site. The plants were then either moved immediately to a room temperature for 2-3 days and imaged, or the temperature was slowly raised to 16°C temperature for 7 days to induce growth before imaging. We used the plants at the vegetative phase of development.

For the imaging in the laser confocal microscope, plants were dissected in filtered sea water using tweezers and fine medical needles under a stereo-microscope (Nikon), so that all leaf primordia covering the SAM were cut off. Isolated shoot apices (SAMs with the youngest leaf primordia) and axillary meristems were fixed and prepared for the imaging according to ClearSee-based clearing method^40^. Isolated apices were fixed with 4% paraformaldehyde (PFA) dissolved in the PBS buffer (pH = 6.9-7.0 adjusted with HCl) for at least 2 h (at the first hour - under vacuum). Apices were washed twice in the PBS buffer for at least 2 min, and incubated for 7-18 days in the ClearSee solution (2 % urea, 10 % xylitol, 15% sodium deoxycholate) at room temperature with a gentle stirring. The ClearSee solution was changed every 1-2 days. Cell walls were stained with 0.05 % Fluorescent Brightener 28 (FB, Sigma) dissolved in the ClearSee solution for at least 30 min, rinsed in the ClearSee solution, and washed in fresh water for 1-2 min.

For the imaging, the apices were mounted in small containers filled with 5% of low-melting point agarose and kept in fresh water. The imaging was performed using an upright confocal laser-scanning microscope (Leica TCS SP8) with long-working distance water-immersion 40x objective. For the FB excitation and emission 405 nm and 425-475 nm wavelengths were used, respectively. Images were collected at 12 bits. Scanning speed was set at 400 Hz with 512 × 512 or 1024 × 1024-pixel frames, the zoom at 0.75-2.0, and z-step at 0.3-0.8 µm. The pinhole was set at 1AE.

### Image Processing and Analysis

Original confocal z-stack images (LIF) were converted in the Fiji (https://fiji.sc) to TIFF files, which were then processed with the MorphoGraphX (MGX)^41^ to obtain top or site views and optical sections. To analyse the structure of apices, a series of optical 2-4 µm thick sections were performed parallel and perpendicular to the SAM major axis (longitudinal and transverse sections, respectively). Developmental stages of leaf primordia were estimated based on optical transverse sections through the apex. The p1 is the youngest primordium apparent as a bulge at the SAM surface. The successive stages were numbered in ascending order (p2, p3, etc., Supplementary Figs. 7-9).

To estimate the number of stem cells at the SAM surface, cell clones were analyzed (Supplementary Fig. 9). Cell clones (usually containing 4-16 cells) were recognized based on the history of cell divisions at projections of SAM anticlinal cell walls. Specifically, the FB signal was projected in the MGX software from the defined depth (0-3 µm) onto the SAM surface. At these projections, the signal is the most intense in newly formed cell walls corresponding to most recent cell divisions (higher order divisions). The signal in the oldest cell walls (regarded as clone borders) is the weakest due to a furrow formed over time between descendant cells.

### Parameterizing the model for eelgrass

The modelling for eelgrass was focused on layer L1. New module formation was implemented by module branching, reflecting the fact that in eelgrass the new shoot apical meristem is not directly derived from the stem cells (Supplementary Note 2). The following parameter range was used: *b* =122 /yr (ref^26^); *r*= 3-8/yr (ref ^29,30^); *N*=7-12; *N*_0_=1-7; *Z*=1000; μ=0.0069. Both symmetric and asymmetric cell division were considered by setting, *λ*=*b*/2, *γ*=0 or *λ*=0, *γ*=*b*, respectively.

### Eelgrass genets of known age cultured in the lab

#### 4-year old eelgrass genets

Three small eelgrass patches, consisting of 17-25 leaf shoots were collected in April 2019 from an eelgrass meadow in Kiel, Germany (Falckenstein, 54.392°N, 10.192°E). To confirm clonal identity each patch was carefully excavated by divers to examine rhizome connections and additionally genotyped with 9 microsatellite loci^42^. In the Baltic Sea, seeds germinate in March or April, while plants become mature at the end of year one. The observed number of shoots can be obtained by branching in the second year. Hence, we infer that the collected eelgrass patches were likely founded by seeds that germinated in 2017, and started branching in 2018. Plants were tagged and transferred to separate plastic boxes in the flow-through seawater system in GEOMAR Helmholtz Center for Ocean Research Kiel, the “Zosteratron”. Leaf shoot number was regularly reduced to allow clones to regrow and branch. In 2022, three years after start of the cultivation, one leaf shoot from each of boxes was selected and resequenced to ~1000x coverage using a Novaseq 6000 S4 platform (paired end reads of 150bp). The time between a collected leaf shoot and the initial mature seedling was four years (3 years in the lab + 1 year in the field). Sequence data are available at BioProject no. PRJNA1025927, accession no. SRR26321801-804 and SRR26321811-812.

#### 17-yr old eelgrass genets

Data are from a whole-genome resequencing of two eelgrass genets with a known age of 17 years^43^. Each genet was initiated by a single shoot collected from Bodega Harbour, California, in July 2004. Before sample collection plants had been kept for 17 years in large, 300-L outdoor flow-through mesocosms at Bodega Marine Laboratory (BML) under ambient light and temperature conditions^44^. Six and five ramets were collected from each genet for genomic analysis in 2021, respectively. The clone assignment was checked based on shared heterozygosity^43^. Ilumina sequencing data are available in the NCBI short read archive (~80x, BioProject no. PRJNA806459, SRA accession nos. SRR18000159–SRR18000170).

#### Sampled eelgrass genets in the field ES_C01-ES_C03

We conducted novel whole-genome resequencing for 10 leaf shoots collected from an eelgrass meadow in Estonia (Supplementary Fig. 14). They were chosen from a larger sampling based on microsatellite data that suggested they belong to 3 genets, containing 3, 4, and 3 ramets, respectively. This was confirmed by whole-genome SNPs. The clonal lineages were named from “ES_C01” to “ES_C03” in this study. Data are available in BioProject no. PRJNA1025927, SRA accession nos. SRR26321797-SRR26321810.

#### YU20_FI

Whole-genome resequencing for 24 ramets of a single large eelgrass genet was conducted in Finland at Ängsö^12^. The next-generation sequencing data are available in the NCBI short read archive (~80x, BioProject no. PRJNA557092, SRA accession nos. SRR9879327-SRR9879353).

### YU23_C01-YU23_C11

In a large population data set encompassing Pacific and Atlantic sites, 190 ramets from 16 geographic locations were re-sequenced^31^, which revealed 11 genets in total that comprised 2-13 ramets. Previously, only one ramet per detected genet was included in subsequent phylogeographic analyses. Here, genets were named “YU23_C01” to “YU23_C11” and their among-ramet genetic differentiation was used for age determination. Next-generation sequencing data are available in the NCBI short read archive^31^.

### Whole-genome resequencing data of new populations

Bulk DNA of the meristematic region and the basal portions of the leaves was extracted using NucleoSpin Plant II kit (Macherey-Nagel, Germany). DNA concentration was determined using a Qubit Fluorometer (Thermo Fisher Scientific) and Nanodrop Spectrophotometer (Thermo Fisher Scientific), and DNA quality was checked by agarose gel electrophoresis. DNA was sent to Beijing Genomics Institute (Hong Kong) for library construction and sequencing. The libraries were sequenced on either Novaseq 6000 S4 platform (PE150bp) or Hiseq Xten platform (PE150bp).

### Mapping the sequencing data to the reference genome

We assessed the quality of the raw reads using FastQC v0.11.7 (https://www.bioinformatics.babraham.ac.uk/projects/fastqc/). BBDuk (https://jgi.doe.gov/data-and-tools/bbtools/bb-tools-user-guide/bbduk-guide/) was used to remove adapters and for quality filtering according to the following criteria (1) sequence downstream with quality < 20 was trimmed (trimq=20); (2) reads shorter than 50 bp after trimming were discarded (minlen=50); (3) reads with average quality below 20 after trimming were discarded (maq=20). FastQC was used to do a second round of quality check for the clean reads. Clean reads were then mapped against the *Z. marina* reference genome v2.1 (ref^45^) using BWA-MEM v0.7.17 (ref^46^) with default parameters. The aligned reads were sorted using SAMtools v1.7 (ref^47^), and duplicated reads were marked using MarkDuplicates tool in GATK v4.0.1.2 (ref^48^). Only properly paired reads (0×2) with MAPQ of at least 20 (-q 20) were kept using SAMtools.

### Clone assignment check for ramets collected from Estonia

GATK4 was used to conduct joint SNP calling for the 10 ramets. HaplotypeCaller was used to generate a GVCF format file for each individual, and GenotypeGVCFs was used for SNP calling based on the combined GVCF file from CombineGVCFs. After filtering (github), the shared heterozygosity method^43^ was used to detect clonemate pairs.

### SNP calling with cancer callers and calculation of VRF50(X1, X2)

Eelgrass (*Zostera marina*) is diploid, and ~99.67% of the genome is homozygous, based on which we assumed that a somatic mutation always changes a homozygous genotype to a heterozygous genotype. The software packages Mutect2 (ref^49^) and Strelka2 (ref^50^) were used for SNP calling. They compared the “normal” sample and the “tumor” sample. Here, SNPs were assumed to represent the ancestral “normal” case if homozygous for the reference allele, because most novel mutations will turn a homozygous to a heterozygous site. Accordingly, the “tumor” sample carried the novel alternative allele. For a specific Mutect2/Strelka2 run with X1 as the “normal” sample and X2 as the “tumor” sample, we used VRF50(X1, X2) to represent the number of somatic mutations in X2 with the variant read frequency ≥ 0.5. VRF50(X1, X2) was calculated as the number of SNPs meeting the following criteria: 1) the coverage of X1 ≥ 12; 2) the coverage of X2 ≥ 23; 3) the variant read frequency of X1 <= 0.01; 4) the variant read frequency of X2 ≥ 0.50.

### Calculation of VRF50(R_x_)

During clonal growth, the fixation of SoGV within all the stem cells leads to substitutions compared with the founder ramet (for the eelgrass case see Supplementary Fig. 1). We defined S(R_x_) to represent the number of the fixed SoGV (i.e., **S**ubstitutions) in the ramet R_x_ compared with the founder seedling/ramet. By definition, the fixed SoGV have an allele frequency of *f*=0.5 under diploidy. Based on sequencing data, allele frequency could be estimated by the variant read frequency (VRF). In the histogram of VRF, the fixed SoGV form a peak at VRF=0.5 (Supplementary Fig. 1). However, for a normal coverage (<100x), mosaic distribution overlaps with the left-hand part of the fixation distribution. Hence, we only focused on the right-hand part of the fixation distribution, and used VRF50(R_x_) as a proxy for S(R_x_), which was the number of the fixed SoGV with a VRF ≥ 0.5.

After a specific time period from the initiation of the clonal lineage, the number of fixed SoGV in a ramet/module R_x_, S(R_x_), is expected to follow a Poisson distribution, S(R_x_)~Poisson(λ). For a given S(R_x_), the VRF has equal probability to be >0.5 or <0.5, and thus VRF50(R_x_) is assumed to follow a binomial distribution, VRF(R_x_)~B(S(R_x_), 0.5). The expectation of VRF50(R_x_) is 0.5 λ.

We used VRF50(R_x_)_obs_ to represent the value of VRF50(R_x_) detected from the sequencing data that sufficiently cover a subset of the reference genome. To obtain VRF50(R_x_)_obs_, the most straightforward logic would be to compare the founder ramet/seedling and the target ramet R_x_. However, the founder did not exist anymore after it had divided into two daughter ramets. Thus, we did an indirect calculation of the VRF50(R_x_)_obs_ (Supplementary Fig. 13). For example, to obtain VRF50(R01)obs, each of the other collected ramets of the same clonal lineage was used as the “normal” sample in SNP calling (Mutect2 or Strelka2), and the maximum value for VRF50(clonemate of R01, R01) was assigned to VRF50(R01)obs. Both Mutect2 and Strelka2 were used to calculate VRF50(R_x_)_obs_ for the clonal lineages with known age, and the results were similar. For the other analyses, we used only Mutect2.

Note that the sequencing data only sufficiently cover a subset of the genome. To estimate the genome coverage, HaplotypeCaller (GATK4) was run for each ramet using BP_RESOLUTION mode (-ERC BP_RESOLUTION). We then counted the number of the nucleotide sites with coverage ≥ 23 (i.e., Size_e). The average VRF50(R_x_) for a clonal lineage was calculated as (average VRF50(R_x_)_obs_)/ (average Size_e) * total genome size. The 95% confidence interval of the average VRF50(R_x_) was estimated based on Poisson distribution, i.e., average VRF50(R_x_) ± 1.96 * sqrt(average VRF50(R_x_)) (Supplementary Data 1).

### Estimating the age of a specific clone

The average VRF50(R_x_) and the age for the clonal lineages with known age were used to fit a linear model (Fig. 4, y = 0.5044x-1.4641, adjusted *R*^2^: 0.9483, p < 0.001), based on which the age of other clones was estimated (Fig. 4, Supplementary Data 1).

## Acknowledgements

This work has been funded by the Human Frontiers in Science (HFSP), grant number RGP_0042_2020 to I.B.B., B.W. and T.B.H.R.. B.W. is also supported by a Barts Charity Lectureship (grant no. MGU045) and a UKRI Future Leaders Fellowship (grant no. MR/V02342X/1). J.K is supported by the Horizon Europe Programme MARBEFES project (grant no. 101060937). We thank Fabian Wendt for maintaining seagrass cultures and Marja Timmermans for providing access to the confocal microscopy.

