## Supplementary Notes and Figures for "Precise age estimation in clonal species using a somatic genetic clock"

### Supplementary Note 1

### Modelling the accumulation of fixed somatic genetic variation in a clonal organism

#### 1.1 Introduction

The model of a clonal organism, represented by a population of modules, is described in the Materials and Methods section of the main text. Here, we expand on the results presented in the main text, considering a range of parameter regimes as well as different methods of maintaining module homeostasis (symmetric vs. asymmetric cell division) and two mechanisms for forming new modules: module splitting and module branching.

We are concerned with the accumulation of fixed somatic genetic variation (SoGV) within modules in the population. The rate of accumulation of fixed neutral SoGV is equal to $\nu$, the mutation rate per cell per unit time (Kimura 1968). This is true for mutations fixating within species or populations of organisms, as well as within modules, and is straightforward to understand. The fixation probability for a SoGV that arises in a single cell within a module of size $N$ is equal to $1/N$. Within the module as a whole, new SoGV arises at a rate $N\nu$. Thus, the accumulation rate of fixed SoGV is $N\nu \times 1/N = \nu$, which is a constant and independent of the module size. This implies that the accumulation of neutral fixed SoGV is linear with time, providing the basis for the somatic genetic clock.

The fixed SoGV accumulation rate does not depend on the time it takes individual SoGV to fixate. However, for the above argument to hold we assume that the timescale in consideration is much larger than the time to fixation (Kimura and Ohta 1971). These fixation times will depend on the model parameters, choice of asymmetric vs. symmetric cell division and choice of module splitting vs. module branching.

In the following we first establish that linear accumulation of fixed SoGV is reached in all model regimes over a long time period, and consider how the rate of fixed mutation accumulation relates to model parameters. We then consider the timescales over which the linear approximation is valid, first by approximating the expected conditional fixation times for the different process within our model that lead to fixation and comparing these with simulation data. Finally, we consider the accumulation of fixed mutations during the early period of population growth and how the time to linearity relates to the conditional fixation times.

#### 1.2 Linear accumulation of fixed somatic genetic varion

In order to verify that neutral mutations accumulate linearly within our model, we have generated time-series simulation data for both symmetric and asymmetric cell division. Results are plotted for $\mu = 1$ and $\mu = 0.01$ in Supplementary Figs. 2 and 3, respectively. We vary the number of stem cells per module $N$, the number of founder stem cells that are recruited to form new modules $N_0$ and the module formation rate $r$. The cell division rate $b = 120/\text{year}$ and population size $Z = 100$ are kept constant. We have limited to module splitting here. However, examples for module branching are shown in Figure 2 of the main text. For almost all parameter regimes we consider, and both types of cell division, fixed SoGV accumulates linearly with time, over a time period of two hundred years. The exception is for asymmetric cell division when $N = 100$, $N_0 = 50$, linearity is not yet reached for smaller values of $r$ (new module formation rate). However, we expect that if we increased the time period this case would also appear linear.

The accumulation rate of fixed SoGV depends on whether symmetric or asymmetric cell division is used, as is evident from Supplementary Figs. 2 and 3. We can estimate the fixed SoGV accumulation rate by estimating $\nu$, the mutation rate per cell per year. For symmetric cell division that occurs at rate $b$, there are $2\mu$ new mutations on average at each cell division and cell divisions occur at a rate $b$. This is true regardless of whether the module is in homeostasis or growth. Thus, we expect that $\nu = 2\mu b$.

For asymmetric cell division it is slightly more complicated, as the expected number of new mutations per cell at each division is $2\mu$ during module growth, but only $\mu$ during homeostasis. We make the approximation $\nu \approx \mu b$. This will underestimate the true fixed mutation accumulation rate, because it ignores the increase in mutations during module growth. However, we expect that this effect will be small, and will decrease with $r$ (new module formation rate), as the growth phase becomes negligible compared to homeostasis.

In Supplementary Fig. 4 we plot the fixed SoGV accumulation rates obtained by least-squares linear regression on the data presented in Supplementary Fig. 2. For symmetric cell division the estimated rate is always close to but slightly less than $2\mu b$. The estimated rate is closer to $2\mu b$ for smaller $r$ and for larger $N_0$. Conversely, for asymmetric division the estimated rate is close to but slightly greater than $\mu b$. It is closer to $\mu b$ for smaller $r$ and for larger $N_0$. In the case of $N = 100, N_0 = 50$, the estimated rate for asymmetric division does go below $\mu b$ for small values of $r$. However, it is clear from Supplementary Fig. 2, that linearity is not reached within two hundred years in this case and therefore, we would expect an underestimate of the true rate.

#### 1.3 Time delay before reaching linearity

We have demonstrated that fixed SoGV accumulates linearly in modules over long time periods. To use the somatic genetic clock to estimate population age, it is necessary that linearity is reached sufficiently quickly, i.e. before the age of the youngest clone used to calibrate the clock (four years for the eelgrass samples we considered in the main text). To obtain a quantitative insight into how the time to linearity depends on the specifics of the model we estimate the conditional fixation times under different regimes. This is the time for SoGV, which arises as a mutation in a single cell, to become fixed within the module, given that it does fix. As we have discussed, the equilibrium accumulation rate of fixed SoGV does not depend on this conditional fixation time. However, the time to reach the equilibrium will increase with increasing fixation times (Kimura and Ohta 1971). We assume that the mean conditional fixation time gives a lower bound on the time to linearity.

We begin by considering the different processes that lead to fixation within our model and approximate the conditional fixation times for each, if taken in isolation. We then compare these to conditional fixation times from the full model by simulating a single module. Finally, we look at simulated data for the number of fixed SoGV in populations of modules over short time periods and consider how the conditional fixation times relate to the time to linearity.

##### 1.3.1 Theoretical predictions for conditional fixation times

In our model SoGV can become fixed either due to cell turnover during module homeostasis or the sampling process that occurs during the formation of new modules. The first process is essentially a Moran process, if cell division is symmetric. If cell division is asymmetric fixation cannot occur during homeostasis as frequencies of SoGV do not change, therefore fixation occurs exclusively due to the second process. The second process, the formation of new modules, we approximate by a modified Wright-Fisher process with different sampling protocols depending on whether new modules are formed by splitting or branching. In the following we estimate the expected conditional fixation time for SoGV arising in a single cell, in a single module, subject to one of these processes.

###### Moran process

The expected conditional fixation time for a single initial mutant in a population of size $N$ subject to a discrete time Moran process has been derived by Antal and Scheuring (2006). For a neutral mutant it is given by

$$\tau_{\text{Moran}} = (N-1)^2 \,, \tag{1.1}$$

assuming time step is size 1. In our model the average time between update events during module homeostasis is $1/Nb$, thus we can write the expected conditional fixation time

$$T_{\text{Moran}}(b) = \frac{(N-1)^2}{Nb} \,. \tag{1.2}$$

For a sufficiently large population this becomes $T_{\text{Moran}}(b) \approx N/b$.

###### Modified Wright-Fisher process

We estimate the expected conditional fixation time due to the successive formation of new modules. We consider a single module that at each time step produces a new child module. The original parent module is then either retained or replaced by the child module, each with 50% probability. This process is approximated as a modified Wright-Fisher model (MWF) with an additional sampling step. The first sampling step corresponds to the formation of a new module and the second sampling step approximates the growth phase of the module. The details of each sampling step depend on whether new modules are formed by branching or splitting:

- Module splitting: A sample of $m$ cells is taken from the parent module *without* replacement to create a pool of founder cells. We let $m = N_0$ or $N - N_0$ each with 50% probability, corresponding to the cases where the child and parent module is retained, respectively. From the founder cells, $N$ cells are sampled *with* replacement, which form the new module after growth.

- Module branching: There is 50% chance that the parent module is retained, in which case no change is made to the module. If instead a new child module is produced, a sample of $N_0$ cells is taken from the parent module *without* replacement to create a pool of founder cells. From these founder cells we sample $N - N_0$ additional cells *with* replacement. These are combined with the founder cells to form the new child module after growth.

We are not aware of an exact solution for the conditional fixation time for this process. However, we can obtain a solution for large module size $N$ by using a diffusion approximation. For a general finite population with neutral selection and discrete time steps, the expected conditional fixation time is (Kimura and Ohta 1969)

$$\tau(p) = \int_p^1 \frac{2x(1-x)}{V_{\delta x}} dx + \frac{1-p}{p} \int_0^p \frac{2x^2}{V_{\delta x}} dx\,, \tag{1.3}$$

where $p$ is the initial proportion of mutants and

$$V_{\delta x} = \mathrm{Var}[\delta X | X = x] \tag{1.4}$$

is the variance of the rate of change in the proportion of mutants per generation. Here, $\delta X = X' - X$, where $X$ is the proportion of mutants at time $t$ and $X'$ the proportion at time $t' = t+1$. As we are assuming neutral selection, the mean rate of change is $\mathrm{E}[\delta X | X = x] = 0$.

To compute $V_{\delta x}$, let $Y = NX$ and $Y' = NX'$, i.e. the number of mutants in the population at time $t$ and and $t'$, respectively. Then,

$$\begin{aligned} N^2 V_{\delta x} &= \mathrm{Var}[\delta Y | X = x] \\ &= \mathrm{E}[(\delta Y)^2 | X = x] - \mathrm{E}[\delta Y | X = x]^2 \\ &= \mathrm{E}[(Y' - Y)^2 | X = x] \\ &= \mathrm{E}[Y'^2 | X = x] - N^2 x^2 \\ &= \mathrm{Var}[Y' | X = x]\,, \end{aligned} \tag{1.5}$$

where we have used the fact that under neutral selection $\mathrm{E}[Y'] = E[Y] = Nx$.

**Module splitting**

First, we consider the case of module splitting. Let $M$ be the total number of founder cells that are sampled without replacement from the module. Then the number of founder cells that are mutants $K$, conditioned on $M = m$ and $X = x$ follows a Hypergeometric distribution, with mean and variance given by

$$\mathrm{E}_H[K | m, x] = mx \tag{1.6}$$

$$\mathrm{Var}_H[K | m, x] = mx(1-x)\left(\frac{N-m}{N-1}\right). \tag{1.7}$$

From the founder cells, $N$ cells are sampled with replacement, thus the resulting number of mutant cells $Y'$, conditioned on $K = k$ and $M = m$, follows a binomial distribution with mean and variance given by

$$\mathrm{E}_B[Y' | k, m] = \frac{Nk}{m} \tag{1.8}$$

$$\mathrm{Var}_B[Y' | k, m] = \frac{Nk}{m}\left(1 - \frac{k}{m}\right). \tag{1.9}$$

The law of total variance gives

$$
\begin{aligned}
\mathrm{Var}_C[Y'|m,x] &= \mathrm{E}_H[\mathrm{Var}_B[Y'|k,m]|m,x] + \mathrm{Var}_H[\mathrm{E}_B[Y'|k,m]|m,x] \\
&= \mathrm{E}_H\left[\frac{NK}{m}\left(1-\frac{K}{m}\right)\middle| m,x\right] + \mathrm{Var}_H\left[\frac{NK}{m}\middle| m,x\right] \\
&= \frac{N}{m^2}\left\{\mathrm{E}_H[K|m,x]\left(m-\mathrm{E}_H[K|m,x]\right) + (N-1)\,\mathrm{Var}_H[K|m,x]\right\} \\
&= \frac{N}{m^2}\left\{m^2x\,(1-x) + (N-1)mx(1-x)\left(\frac{N-m}{N-1}\right)\right\} \\
&= \frac{N^2x(1-x)}{m}\,. \qquad (1.10)
\end{aligned}
$$

Now we define a probability distribution

$$
P(M=m) = \begin{cases} 1/2 & \text{if } m \in \{N_0,\, N-N_0\}, \\ 0 & \text{otherwise.} \end{cases} \qquad (1.11)
$$

Then from Equation (1.5)

$$
\begin{aligned}
V_{\delta x}^{\text{split}} &= \frac{1}{N^2}\,\mathrm{Var}[Y'|x] \\
&= \frac{1}{2N^2}\left(\mathrm{Var}_C[Y'|N_0,x] + \mathrm{Var}_C[Y'|(N-N_0),x]\right) \\
&= \frac{Nx(1-x)}{2N_0(N-N_0)}\,. \qquad (1.12)
\end{aligned}
$$

**Module branching**

For the case of module branching, the number of mutants after both sampling steps is $K+L$, where $K$ is the number of mutants out of the $N_0$ cells drawn in the first sampling step and $L$ is the number of mutants out of $N-N_0$ cells drawn in the second sampling step. The distribution of $K$ is the same as for module splitting, except that $M$ is not a random variable. Thus, $\mathrm{E}_H[K|x]$ and $\mathrm{Var}_H[K|x]$ are given by Equations (1.6) and (1.7), respectively, setting $m = N_0$. The random variable $L$ conditioned on $K=k$ is binomially distributed with mean and variance given by

$$
\mathrm{E}_B[L|k] = (N-N_0)\left(\frac{k}{N_0}\right) \qquad (1.13)
$$

$$
\mathrm{Var}_B[L|k] = (N-N_0)\left(\frac{k}{N_0}\right)\left(1-\frac{k}{N_0}\right)\,. \qquad (1.14)
$$

The module is only updated with 50% probability at each time step (i.e. if the child module is retained) and thus, from Equation (1.5) have

$$
\begin{aligned}
V_{\delta x}^{\text{branch}} &= \frac{1}{N^2}\,\mathrm{Var}[Y'|x] \\
&= \frac{1}{2N^2}\,\mathrm{Var}[K+L|x] \\
&= \frac{1}{2N^2}\left\{\mathrm{Var}[K|x] + \mathrm{Var}[L|x] + 2\,\mathrm{Cov}[L,K|x]\right\}\,. \qquad (1.15)
\end{aligned}
$$

The first term $\mathrm{Var}[K|x] = \mathrm{Var}_H[K|N_0, x]$ is given by Equation (1.6). The second term is computed in the same way as Equation (1.10), using the law of total variance:

$$\mathrm{Var}[L|x] = \mathrm{E}_H[\mathrm{Var}_B[L|k]||x] + \mathrm{Var}_H[\mathrm{E}_B[L|k]|x] \tag{1.16}$$
$$= \left(\frac{N - N_0}{N_0}\right)\left(N - \frac{N_0(N - N_0)}{N - 1}\right) x(1 - x)\,. \tag{1.17}$$

The third term is given by

$$\mathrm{Cov}[L, K|x] = \mathrm{E}[LK|x] - \mathrm{E}[L|x]\,\mathrm{E}[K|x]\,. \tag{1.18}$$

We know $E[K|x]$ is given by Equation (1.6) and

$$\mathrm{E}[L|x] = \mathrm{E}_H[E_B[L|k]|x] = (N - m)x\,, \tag{1.19}$$

where we have substituted in Equations (1.6) and (1.13). Finally we compute

$$\begin{aligned}\mathrm{E}[LK|x] &= \sum_{l,k} lk \cdot \mathrm{P}(L = l, K = k|X = x) = \sum_{l,k} lk \cdot p_B(l|k)p_H(k|x)\\ &= \sum_k kp_H(k|x)\,\mathrm{E}_B[L|k] = \frac{N - m}{m}\sum_k k^2 p_H(k|x)\\ &= \frac{N - m}{m}(\mathrm{Var}_H[K|x] + \mathrm{E}_H[K|x]^2)\\ &= \frac{(N - m)^2}{N - 1}x(1 - x) + m^2x^2\,.\end{aligned} \tag{1.20}$$

Again, we have used Equations (1.6), (1.7) and (1.13). Thus, Equation (1.18) becomes

$$\mathrm{Cov}[L, K|x] = \frac{(N - m)^2}{N - 1}x(1 - x)\,. \tag{1.21}$$

Substituting Equations (1.7), (1.17) and (1.21) into Equation (1.15) we get

$$V_{\delta x}^{\mathrm{branch}} = \frac{(N - N_0)(N - 1 + N_0)}{2NN_0(N - 1)}x(1 - x)\,. \tag{1.22}$$

**Conditional fixation times**

We can write $V_{\delta x} = x(1 - x)/N_e$, where $N_e$ is an effective population size that depends on the model. Substituting into Equation (1.3), we obtain an expression for the expected conditional fixation time:

$$\tau_{\mathrm{MWF}}(p) = -\frac{1}{p}2N_e(1 - p)\log(1 - p)\,, \tag{1.23}$$

assuming time step is size 1. For time step $\delta t$ and taking the limit $p \to 0$, we obtain an approximation for the conditional fixation time for a single initial mutant:

$$T_{\mathrm{MWF}} = 2N_e\delta t\,. \tag{1.24}$$

Using Equations (1.12) and (1.22) we can write down the effective population sizes for module splitting and module branching, respectively:

$$N_e^{\mathrm{split}} = 2N_0\left(1 - \frac{N_0}{N}\right) \tag{1.25}$$

$$N_e^{\mathrm{branch}} = \frac{2N_0}{\left(1 - \frac{N_0}{N}\right)\left(1 + \frac{N_0}{N-1}\right)} \approx \frac{2N_0}{1 - (N_0/N)^2}\,. \tag{1.26}$$

The approximation in the second equation holds for $N \gg 1$. We can also compare with the standard Wright-Fisher (WF) process, for which $N_e = N$ (Kimura and Ohta 1969). The MWF process with module splitting always has a smaller effective population size compared to the WF process. However, for the MWF with module branching, the effective population size is smaller than that of the WF process only if $N_0 < N/(1+\sqrt{2})$. Smaller effective population sizes correspond to faster fixation.

In our model the time step can be approximated as a sum of the time that a module spends growing and the time it spends in homeostasis, i.e. $\delta t = \delta t_g + \delta t_h$. The expected time in homeostasis is given by $\delta t_h = 1/r$. We approximate the growth time by

$$\delta t_g^{\text{split}} = \frac{1}{2b}\left[\log(N/N_0) + \log(N/(N-N_0))\right], \text{ or} \tag{1.27}$$

$$\delta t_g^{\text{branch}} = \frac{1}{2b}\log(N/N_0), \tag{1.28}$$

for module splitting and module branching, respectively. Thus, the time step is $\delta t = 1/r_{\text{eff}}$, where $r_{\text{eff}}$ is the effective branch rate, given by

$$r_{\text{eff}}^{\text{split/branch}} = \left[\frac{1}{r} + \delta t_g^{\text{split/branch}}\right]^{-1}. \tag{1.29}$$

Note that if $r \ll b$ we can set $r_{\text{eff}} \approx r$. The conditional fixation times for module splitting and module branching are therefore given by

$$T_{\text{MWF–split}}(r_{\text{eff}}^{\text{split}}) = \frac{4N_0}{r_{\text{eff}}^{\text{split}}}\left(1 - \frac{N_0}{N}\right) \tag{1.30}$$

$$T_{\text{MWF–branch}}(r_{\text{eff}}^{\text{branch}}) = \frac{4N_0}{r_{\text{eff}}^{\text{branch}}}\left[\left(1 - \frac{N_0}{N}\right)\left(1 + \frac{N_0}{N-1}\right)\right]^{-1} \tag{1.31}$$

$$\approx \frac{4N_0}{r_{\text{eff}}^{\text{branch}}}\left[1 - (N_0/N)^2\right]^{-1}. \tag{1.32}$$

#### Comparing conditional fixation times under different processes

In Supplementary Fig. 5a we compare the mean conditional fixation times that we have estimated for the Moran process ($T_{\text{Moran}}$), given by Equation (1.2), and for the MWF process with module splitting ($T_{\text{MWF–split}}$) or module branching ($T_{\text{MWF–branch}}$), given by Equations (1.30) and (1.32). We show results for $r = 2.5$/year and $r = 20$/year, with $b = 120$/year and $N = 100$. For the MWF processes we plot the conditional fixation times calculated using both $\delta t = 1/r$ and $\delta t = 1/r_{\text{eff}}^{\text{split/branch}}$. We note that there is a small correction when using the latter, but this is negligible for smaller $r$.

In most cases, mutations fixate faster under the Moran process compared to both MWF processes. This is because we enforce that $b > r$, i.e. cell division occurs faster than the formation of new modules. However, for smaller values of $N_0$, the effective population size may be sufficiently reduced under the MWF that fixation happens faster compared to the Moran process, as can be seen for $r = 20$ in Supplementary Fig. 5a. Comparing Equations (1.2), (1.30) and (1.32), we can see that the conditional fixation times are longer for the Moran process compared to the MWF if

$$\frac{b}{r_{\text{eff}}^{\text{split}}} < \frac{N}{4N_0} \qquad \text{or} \qquad \frac{b}{r_{\text{eff}}^{\text{branch}}} \lesssim \frac{N}{4N_0}\left(1 - \frac{N_0^2}{N^2}\right), \tag{1.33}$$

for module branching and module splitting, respectively. The approximation holds for $N \gg 1$. For module splitting the conditional fixation times are symmetric, and thus, there is an equivalent threshold for large $N_0$. Comparing the two MWF processes, it is clear from Equations (1.30) and (1.32) and Supplementary Fig. 5a that $T_{\text{MWF–split}} < T_{\text{MWF–branch}}$. Thus, fixation is faster for module splitting compared to module branching.

##### 1.3.2 Comparison with simulated conditional fixation times

We ran simulations of our model with only a single module, such that whenever a new module forms either the parent or child module is subsequently removed with equal probability (i.e. $Z = 1$). Setting $r = 8/\text{year}$, $\mu = 0.01$ and $b = 120/\text{year}$ we computed the conditional fixation times for all mutations that fixed within two hundred (symmetric cell division) or one thousand (asymmetric cell division) years, for twenty independent simulations. These results are plotted in Supplementary Fig. 5b for three module sizes: $N = 5$, 20 and 100.

For asymmetric cell division, only the process of new module formation leads to fixation. Thus, we need only compare the simulation data with the MWF predictions. For both module branching and module splitting, the simulated conditional fixation times are close to, but slightly below, the predicted values. This could be due to the assumption that the growth phase is represented by binomial sampling in the MWF model. The true distribution will certainly differ, as the growth phase is not accurately represented by sampling with replacement. Rather, after sampling we would replace both the original cell and an additional copy, before re-sampling. Another reason for deviating from the prediction is that we have assumed that all new mutations occur at frequency $1/N$. However, in the simulated model, some mutations will occur at higher frequency, because they arise during the growth phase when the module size is smaller. Finally, we note that the theoretical predictions are derived under the assumption of large $N$. Although the predicted values slightly overestimate the simulation data, they clearly capture the qualitative behaviour and provide us with a simple method for estimating the fixation times. This is true even for small module size, $N = 5$.

Finally, we consider symmetric cell division and module splitting. In this case, we must consider the affect of the Moran process in addition to the MWF. Thus, we plot the predicted conditional fixation times for both in Supplementary Fig. 5d along with the simulation data. For intermediate values of $N_0$, near $N/2$, $T_{\text{Moran}}$ approximates the simulated conditional fixation times well. In this region, $T_{\text{MWF–split}}$ is very large and so we would expect that most fixation is occurring due to the Moran process. However, as $T_{\text{MWF–split}}$ becomes smaller, we can see that the effect of the combined processes is to cause the resultant conditional fixation times in the simulated model to be smaller than either process would produce in isolation. This suggests that the two processes have a cumulative effect and that the simulated conditional fixation times must be less than $T_{\text{MWF–split}}$ and $T_{\text{Moran}}$. Thus, the smaller of these can be considered an upper bound.

##### 1.3.3 Effect of conditional fixation times on the time to reach an equilibrium rate of fixed mutation accumulation

To consider how the conditional fixation times relate to the time it takes to reach an equilibrium constant rate of fixed mutation accumulation, we plot fixed SoGV against time over a short initial time period (two and four years for symmetric and asymmetric cell division, respectively) in Supplementary Fig. 6. Along with the simulated data we show the expected conditional fixation times calculated for the Moran process from Equation (1.2) (only for the case of symmetric cell division) and for the MWF with module splitting (for both symmetric and asymmetric cell division). Where the conditional fixation times exceed the time limits shown they are stated.

From Supplementary Fig. 6 we can see that longer predicted conditional fixation times correspond to longer periods of non-linearity. For larger values of $N$ and $N_0$ there is a significant lag-phase during which no fixed SoGV is observed. This is particularly clear in the case of $N = 100$, $N_0 = 50$ and for asymmetric cell division. Indeed for $r = 2.5$ there is no fixed SoGV within the four year time period. This is consistent with the fact that $T_{\text{MWF–split}} = 40.6$ years. If we compare the same parameter regime, but for symmetric cell division we see that the plateau is significantly reduced. Although $T_{\text{MWF–split}}$ is the same as in the asymmetric case, $T_{\text{Moran}}$ is significantly lower, thus speeding up fixation time and the progression to linearity.

We can also see that linearity is approached faster for smaller values of $N_0$. Again, this is predicted by the conditional fixation times which decrease with decreasing $N_0$ (see **??**. This effect is clear for the case of asymmetric cell division if we compare the top and bottom rows in Supplementary Fig. 6b. For symmetric cell division the picture is slightly more complicated as we must also take into account the Moran process.

For smaller values of $r$, the Moran process occurs on a much faster time scale than the MWF and has correspondingly smaller conditional fixation times. Thus, we can see in Supplementary Fig. 6a for $r = 2.5$/year, that there is only a weak effect from reducing $N_0$. By contrast, for larger values of $r$ the time scales become closer. For the case of $r = 20$/year and $N = 100$, reducing from $N_0 = 50$ to $N = 1$ means that mutations fixate faster in the MWF compared to the Moran, i.e. $T_{\text{MWF–split}} < T_{\text{Moran}}$. In this case it is clear that the progression to linearity is sped up by reducing $N_0$.

#### 1.4 Application to the somatic genetic clock

The somatic genetic clock relies on the assumption that fixed SoGV accumulates linearly within modules, and thus it can only be utilised for estimating the age of clonal species if linearity is reached within biologically relevant time scales. Calculating the conditional fixation probabilities does not give us a strict rule for the amount of time needed to wait for linearity. However, it does allow us to compare between different model regimes and gives us a minimum bound for the amount of time we need to wait before we can expect the somatic genetic clock to be applicable. Our results suggest that for eelgrass and similar plants that have only a few stem cells, we can reasonably expect the somatic genetic clock to work over time scales of a few years. Even considering that asymmetric cell division and module branching slow down fixation compared to symmetric division and module splitting. However, in species with larger stem cell pools and crucially, if the number of stem cells that initialise new modules is large, fixation times may be too long to apply the somatic genetic clock.

### Supplementary Note 2

### Architecture of the shoot apical meristem of eelgrass (*Zostera marina*)

A somatic mutation can be fixed and propagate infinitely when it occurs in stem cells. In plants, these pluripotent cells are stored at the shoot apical meristem (SAM) and provide cells for the formation of above-ground organs such as leaves or lateral branches. As the fate of somatic mutations depends on the SAM structure, the number and dynamics of stem cells (Klekowski 1988; Burian 2021), we examined the SAM in *Zostera marina* using 3D imaging performed with the laser confocal microscopy.

#### 2.1 The structure of the shoot apical meristem

A mutation is fixed in a plant body, when it is propagated to all tissues including both outer and inner cells. For example, a mutation is a stem cell at the SAM surface can spread to inner tissues via periclinal cell divisions (parallel to the SAM surface) and subsequent displacement of descendent cells. However, the SAM in angiosperms usually shows layered (or stratified) tunica-corpus structure meaning that cell displacement is restricted in one or more outer tunica layers due to prevalence of anticlinal cell divisions (perpendicular to the SAM surface) (Esau 1965; Lyndon 1998). Consequently, such plants develop as chimeras, in which either entire cell layer (periclinal chimera) or only a clonal sector within a cell layer (mericlinal chimera) is genetically distinct (Klekowski 1988; Frank and Chitwood 2016).

Analyzing 24 plants, we found that the SAM in *Z. Marina* consists of one-layered tunica (L1), where cells divide only in the anticlinal direction, and underlying corpus (L2), where cell division plane is not restricted (Supplementary Fig. 7a). This indicates that cell lineages in the L1 and L2 are separated at the SAM and they contain their own sets of stem cells. Nonetheless, it cannot be excluded that periclinal cell divisions do occur occasionally in the L1 disrupting the layered SAM structure. For example, rare L1 periclinal cell divisions with a very low frequency (0.4 %) have been reported for some angiosperms (Stewart and Dermen 1970; Lyndon 1998).

#### 2.2 The formation of axillary meristems

New ramets in *Z. Marina* develop from axillary meristems which are formed in the axil of each leaf (Supplementary Fig. 7b–e). Because their position in the axil, axillary meristems are likely descendants of 3 or 4 stem cells at the SAM (see Section 2.4: Stem cells at the shoot apical meristem). Axillary meristems are initiated in the acropetal sequence after the leaf initiation meaning that the youngest stages of axillary meristems are found at the most apical shoot portion. In particular, the initial bulging leading to the establishment of axillary meristems occurs at the base of the internode in the p3 leaf axil (Supplementary Fig. 7b). The bulging is associated with periclinal cell divisions in the L2, but not in the L1, where only anticlinal divisions occur. Thus, axillary meristems derive from separated L1 and L2 cell lineages originating from the SAM. Also during further development, the structure of axillary meristems resembles the SAM, and periclinal divisions in the L1 are only associated with the formation of leaf primordia (see Section 2.3: The formation of leaves) (Supplementary Fig. 7c–e). The separation of tunica and corpus cell lineages observed in *Z. Marina* during the formation of axillary meristems is common as well in grasses and other angiosperms (Sharman 1945; Esau 1965; Tian and Marcotrigiano 1994).

#### 2.3 The formation of leaves

Although the layered structure is maintained at the SAM and axillary meristems, the separation of the L1 and L2 is disrupted during leaf formation. As in other monocots, early stages of leaf initiation in *Z. Marina* are associated with periclinal cell divisions in the L1 that mark the tip of bulging primordia (Supplementary Fig. 8a) (Sharman 1945; Thielke 1951; O'Connor et al. 2014). Initially, the periclinal divisions occur in few cells, but soon after they are observed in more cells, not only at the primordium tip, but also at adaxial and abaxial sides of leaf primordia (Supplementary Fig. 8b). Consequently, in addition to epidermis, the L1 can give rise to some or most of the internal leaf tissues. This is in agreement with previous observations showing that in monocots a large portion of leaves or even entire leaf derive from the L1 (Sharman 1945; Thielke 1951; Tilney-Bassett 1963; Esau 1965; Stewart and Dermen 1979; Döring et al. 1999)).

#### 2.4 Stem cells at the shoot apical meristem

In most studied angiosperms, the SAM is dome-shaped with 3–4 surface stem cells positioned centrally (Green, Havelange, and Bernier 1991; Zagórska-Marek and Turzanska 2000; Burian, Barbier De Reuille, and Kuhlemeier 2016; Conway and Drinnan 2017). However, the SAM shape in *Z. Marina* is unusual. Namely, it is elliptical (broader in one direction), and flatted along the major axis at the distal region (Supplementary Fig. 7a). This elliptical shape is likely maintained during development, because new organs are formed at SAM flanks parallel to the major axis in alternate (distichous) phyllotactic pattern.

To estimate the number of stem cells, cell clones were analysed at the SAM surface based on cell division history (Material and Methods; Supplementary Fig. 9a–b; Gola and Jernstedt 2011). The position and number of clones ranging from 7 to 12 reflects the number of stem cells (Supplementary Fig. 9c). Accordingly, stem cells are likely arranged along the SAM major axis and their higher number (in comparison to other angiosperms) appears to be related with a unique SAM shape.

Stem cells are generally defined by their ability to generate the cells which retain stem cell function and the cells which enter the differentiation pathway (differentiating cells) (Laux 2003; Heidstra and Sabatini 2014). Specifically, a stem cell can divide asymmetrically generating one descendent stem cell and one differentiating cell, or symmetrically, where either two stem cells or two differentiating cells are generated (Morrison and Kimble 2006).

Asymmetric divisions are associated with a stable configuration of permanently functioning stem cells, while symmetric divisions allow for a stem cell turnover. In most of vascular plants, stem cells are not permanent meaning that although the same set of stem cells can function for some time, eventually a stem cell can be displaced from the SAM center and replaced by a neighbouring stem cell dividing symmetrically (Zagórska-Marek and Turzanska 2000; Gola and Jernstedt 2011; Conway and Drinnan 2017). Thus, the position of stem cells at the SAM defines their function.

There is a relation between the direction of cell growth and the orientation of cell division plane: plant cells usually divide perpendicular to the direction of maximal growth rate (Hejnowicz and Romberger 1984). Thus, cell displacement due to growth can be deduced from the orientation of cell divisions. In *Z. Marina*, cell divisions parallel to the SAM major axis (Supplementary Fig. 9b, full arrowhead) suggest that stem cells divide asymmetrically: the distal descendant cell retains its position and stem cell function, while the proximal cell is displaced towards SAM flanks and ultimately differentiates. However, stem cells can also divide perpendicular to the major axis (Supplementary Fig. 9b, empty arrowhead) that may reflect symmetric cell divisions leading to the establishment of two descendant stem cells. Consequently, this might lead to an increase in the number of stem cells, the SAM expansion along the major axis and increased SAM size. Alternatively, the number of stem cells can remain unchanged due to displacement and resulting elimination of peripheral stem cells.

# 3

### Supplementary Figures

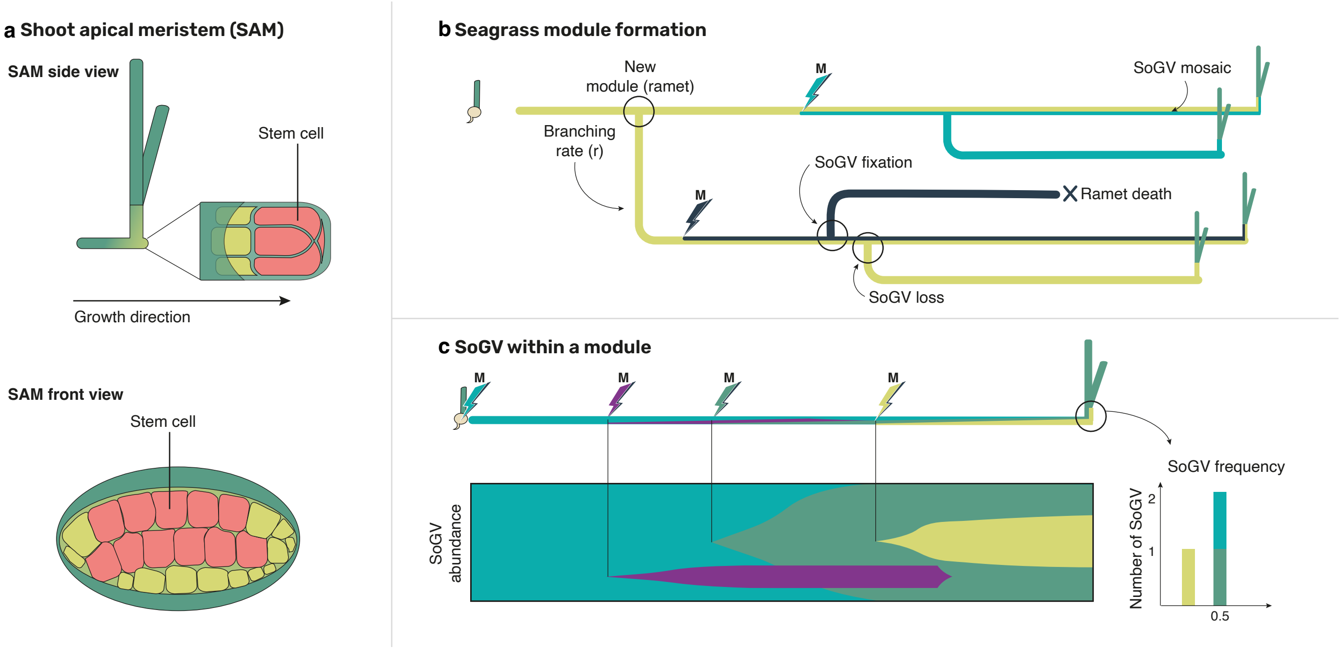

**Supplementary Figure 1: Dynamics of somatic genetic variation in eelgrass (*Zostera marina*). a**, Diagrammatic depiction of the shoot apical meristem, the stem cell population of eelgrass that gives rise to new modules (based on Supplementary Figs. 2–4). **b**, An eelgrass genet is initiated by a single seedling forming the first ramet. Somatic mutations continuously occur during clone branching in space and time, while new SoGV can become fixed or disappear again. c, Owing to somatic genetic drift, somatic mutations that are initially emerging in a mosaic state at frequency $1/N$ ($N$ = number of stem cells) will eventually be either lost or fixed. During their transition from emergence to fixation, they will appear in variant read frequency (VRF) diagrams as modes with $1/N < f < 0.5$, as eelgrass is diploid, and most mutations ($> 99\%$) will involve a transition from homozygous to heterozygous sites.

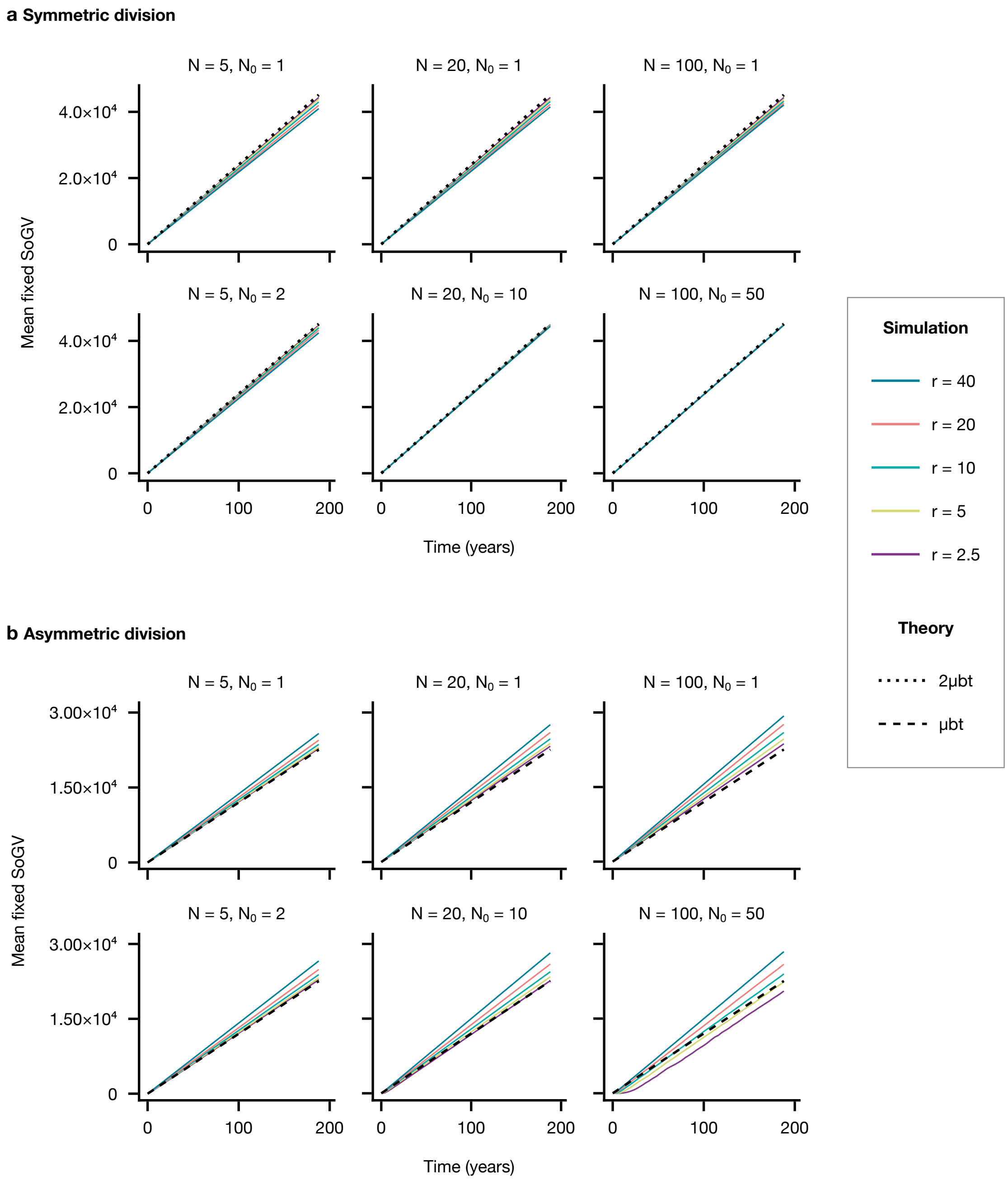

**Supplementary Figure 2: Linear accumulation of fixed SoGV in modules over a long time period** (mutation rate $\mu = 1$). Simulations are shown for a range of parameter regimes, with new modules formed by module splitting and either **a,** symmetric or **b,** asymmetric cell division. At each time point the mean is taken over all modules in twenty simulated populations. *Other parameters:* $Z = 100$ and $b = 120/\text{year}$.

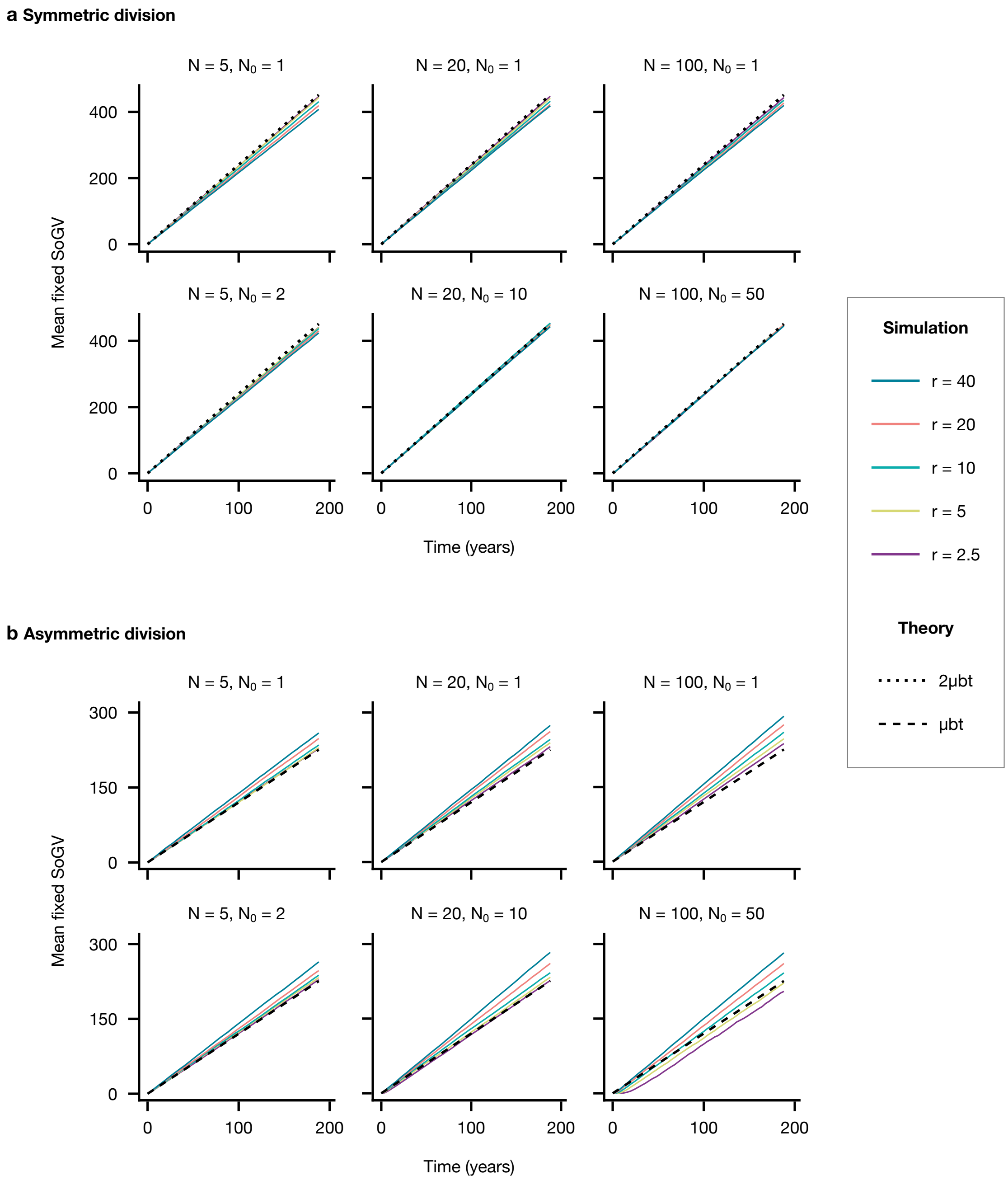

**Supplementary Figure 3: Linear accumulation of fixed SoGV in modules over a long time period** (mutation rate $\mu = 0.01$). Simulations are shown for a range of parameter regimes, with new modules formed by module splitting and either **a,** symmetric or **b,** asymmetric cell division. At each time point the mean is taken over all modules in twenty simulated populations. *Other parameters:* $Z = 100$ and $b = 120/\text{year}$.

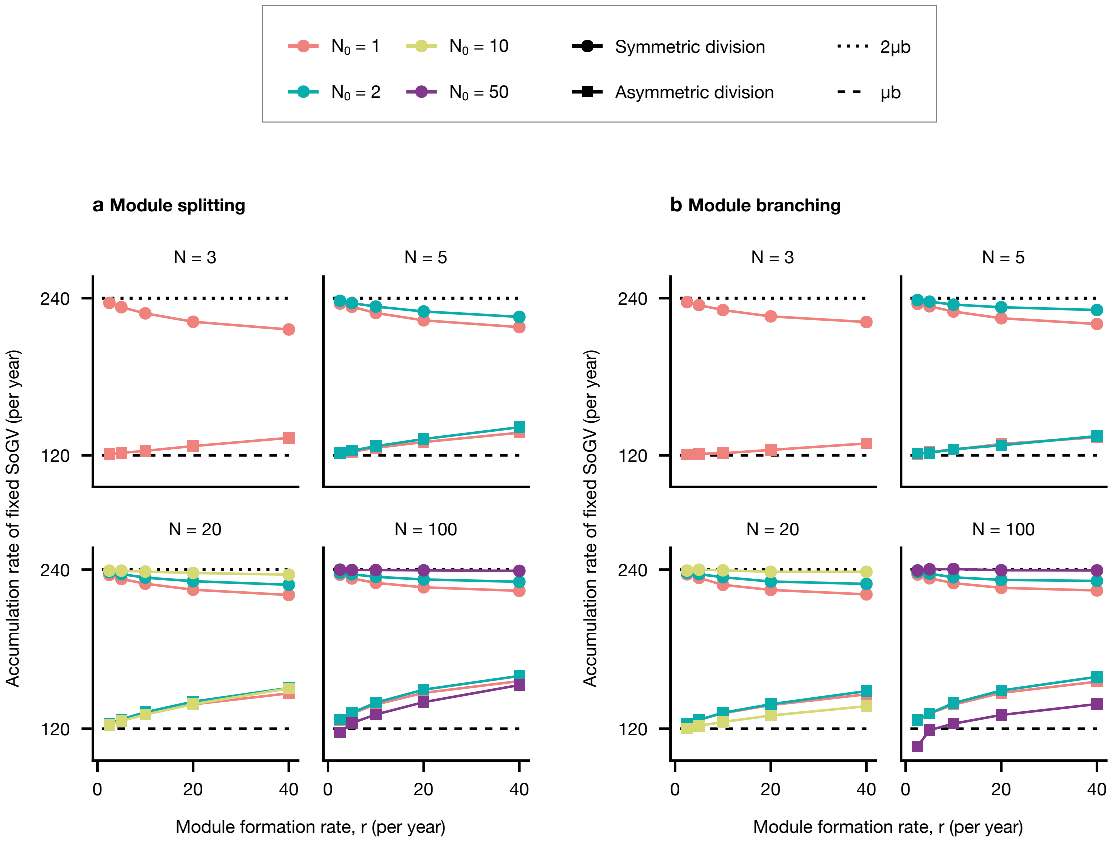

**Supplementary Figure 4: Accumulation rate for fixed SoGV in a population of modules.** Each point is obtained by least-squares regression on data from twenty simulated populations over two hundred years for **a**, module splitting (using data from Supplementary Fig. 2) and **b**, module branching. *Parameters:* $\mu = 1$, $Z = 100$, $b = 120/\text{year}$.

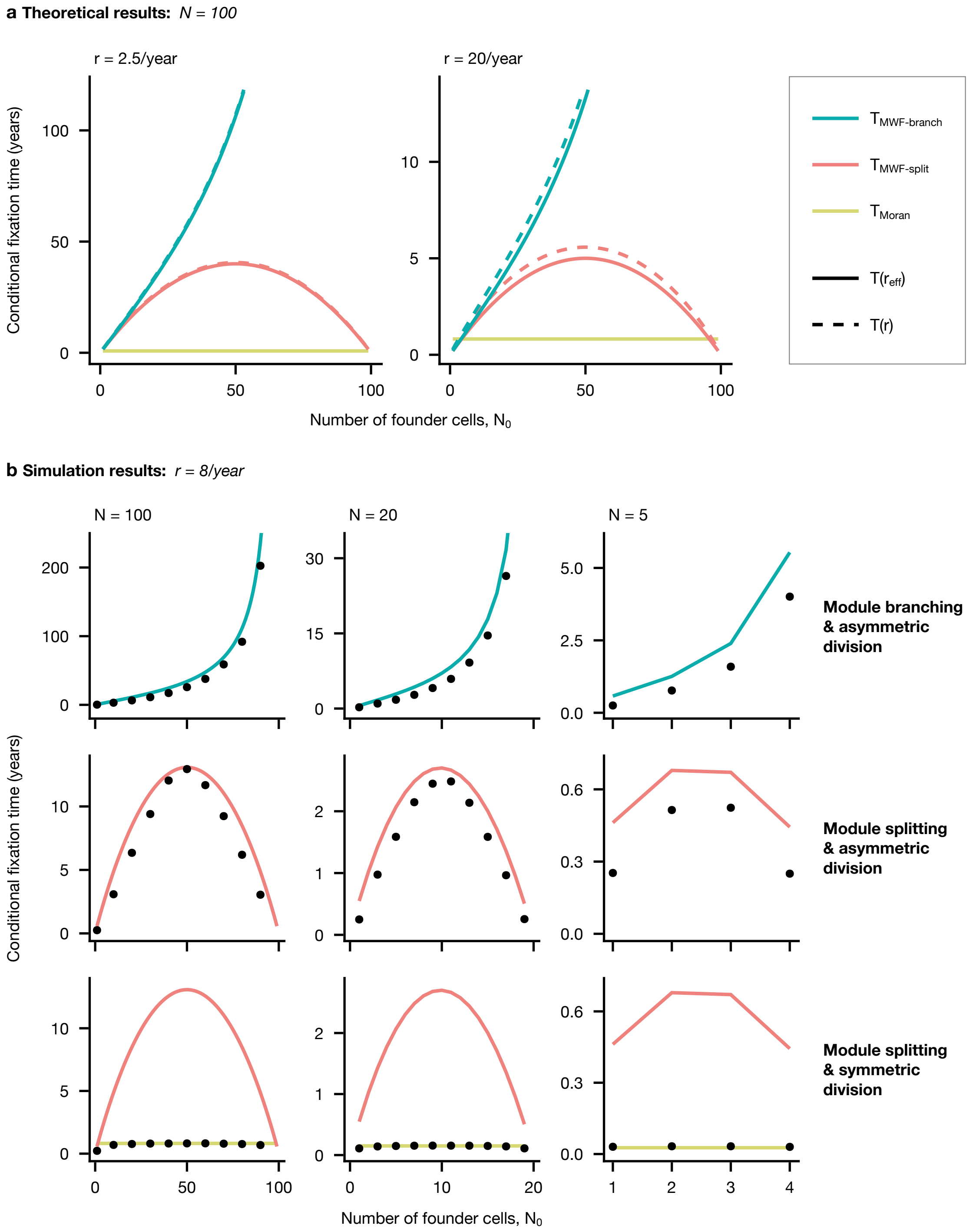

**Supplementary Figure 5: Expected conditional fixation times due to somatic genetic drift within a module. a,** Theoretical predictions for the modified Wright-Fisher (MFW) process with module branching ($T_{\mathrm{MWF-branch}}$), the MFW with module splitting ($T_{\mathrm{MWF-split}}$) and the Moran process ($T_{\mathrm{Moran}}$). **b,** Simulation data for the expected conditional fixation time in a population formed by a single module, calculated by taking the mean over the fixation times for all mutations that become fixed in twenty simulations each run for one thousand (asymmetric division) or two hundred (symmetric division) years. These are compared to the relevant theoretical predictions. *Parameters:* $Z = 1$, $\mu = 0.01$, $b = 120/\mathrm{year}$.

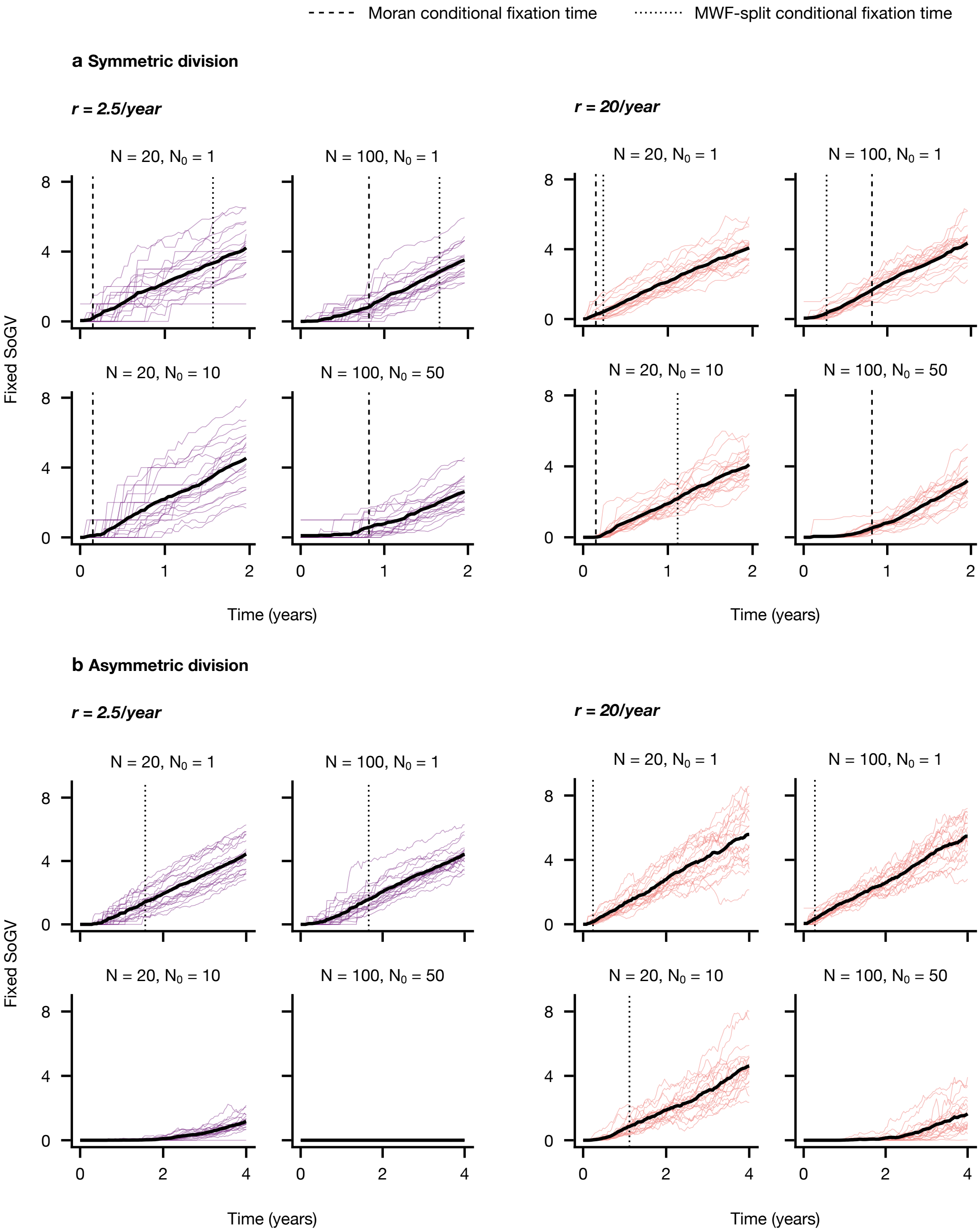

**Supplementary Figure 6: Duration of lag-phase before linear accumulation of fixed SoGV increases with conditional fixation times.** Homeostasis is maintained by either **a,** symmetric cell division or **b,** asymmetric cell division. Thin coloured lines indicate the mean over all modules in a population. Thick black lines are the mean over twenty simulated populations. Dashed lines indicate $T_{\mathrm{Moran}}(b)$ (shown for symmetric cell division only) and dotted lines indicate $T_{\mathrm{MWF\text{-}split}}(r_{\mathrm{eff}})$. For ($r = 2.5$, $N = 20$, $N_0 = 10$), ($r = 2.5$, $N = 100$, $N_0 = 50$) and ($r = 20$, $N = 100$, $N_0 = 50$) $T_{\mathrm{MWF\text{-}split}}(r_{\mathrm{eff}})$ exceeds the time limits shown on the plot and is given by 8.1, 40.6 and 5.6 years, respectively. *Parameters:* $\mu = 0.01$, $Z = 100$, $b = 120/\mathrm{year}$.

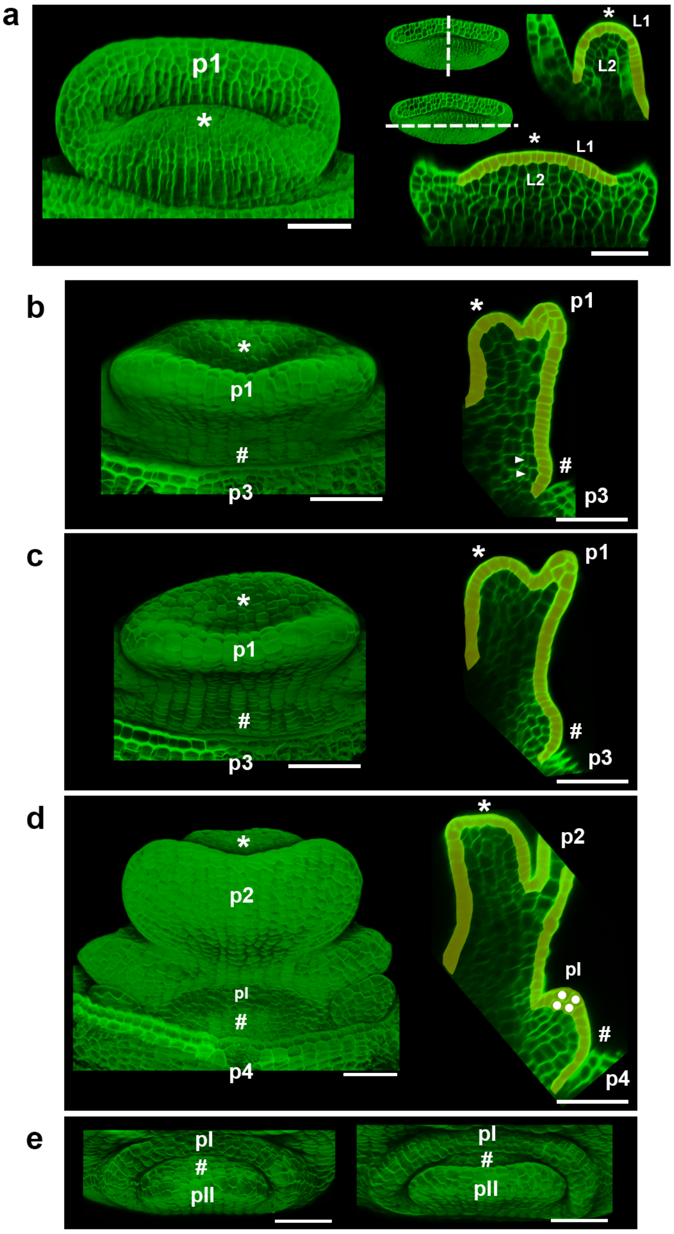

**Supplementary Figure 7: The structure of SAM and axillary meristems in *Zostera marina*.** **a**, The SAM has tunica (L1) – corpus (L2) structure. Left, the site view of the SAM (asterisks); right, optical transverse and longitudinal sections through the SAM. The plane of sections is indicated with dashed lines (inserts). **b–d**, The L1–L2 structure is maintained during the formation of axillary meristems. Left, site views of the shoot apex where consecutive stages of axillary meristem (hash) formation are shown from the initial and bulging (**b**, **c**) until the generation of the first leaf primordium (**d**). Right, corresponding optical transverse sections through the apex. **e**, Top views of axillary meristems at later stages show development of the first and second leaf primordia. The L1 is coloured in yellow. The L2 and L1 cells after periclinal cell divisions associated with formation of the axillary meristem and its primordia are indicated by arrowheads and dots, respectively. Leaf primordia generated at the SAM (p1–4) or at axillary meristems (pI–pII) are numbered from the youngest. Representative examples are shown from n=23 analysed apices contained 2-4 axillary meristems per apex. Scale bars, 50 µm.

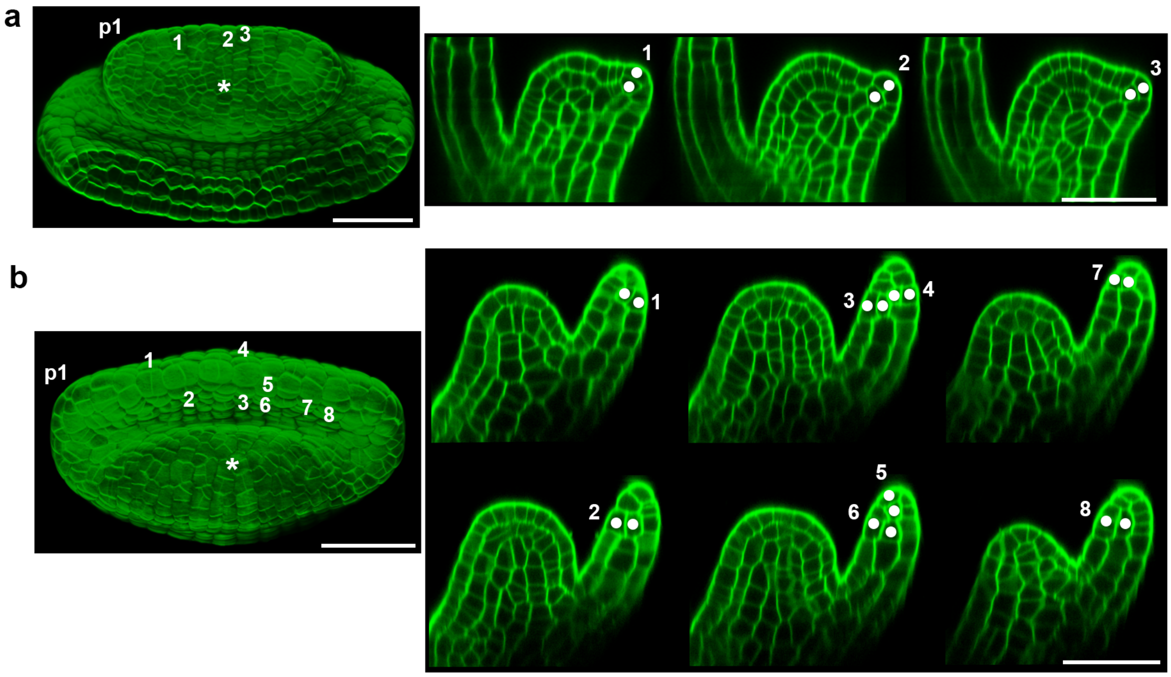

**Supplementary Figure 8: The formation of leaves at the SAM. a,b**, The bulging of p1 leaf primordium (**a**) and its subsequent growth (**b**) at the SAM (asterisks) is associated with periclinal cell divisions in the L1. Left, top views of the apex; right, optical transverse sections through the apex. The L1 cells after periclinal cell divisions are numbered (1–8) and indicated by dots at sections. Representative examples are shown from n=22 analysed leaf primordia at different stages. Scale bars, 50 µm.

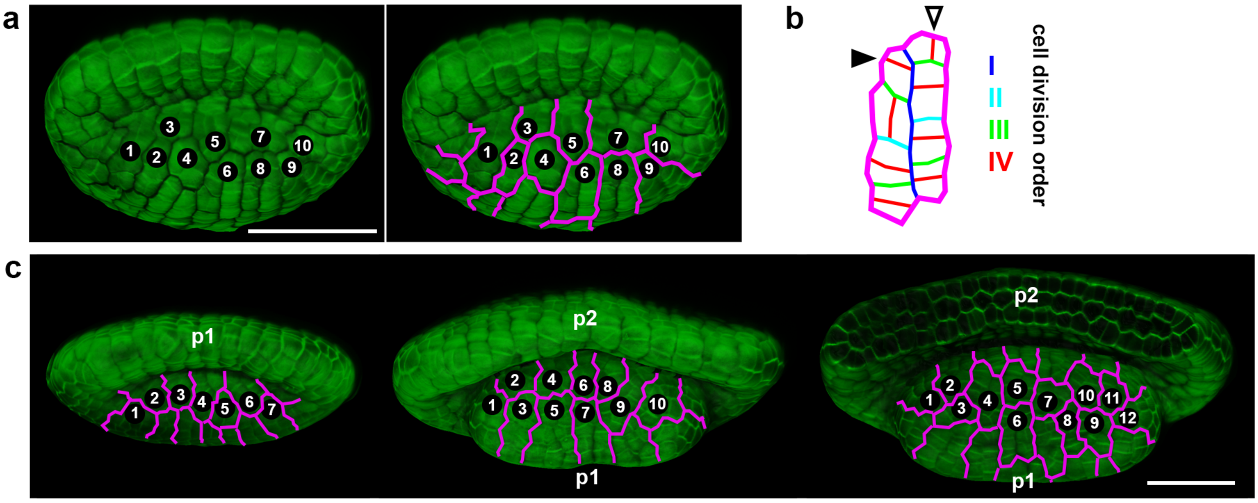

**Supplementary Figure 9: Stem cells at the SAM. a**, The number of stem cells is estimated based on cell clones positioned along the major SAM axis. **b**, An exemplary cell clone (the number 6 at a) is shown with cell division history. The order of cell divisions is indicated with colours: the oldest (blue) and the most recent (red) divisions. Cell divisions parallel (full arrowhead) and perpendicular (empty arrowhead) to the SAM major axis corresponds to asymmetrical and symmetrical stem cell division modes, respectively. **c**, The number of cell clones along the SAM major axis ranges from 7 to 12. The SAMs at different stages of development are shown: left, before the initiation of a new primordium; middle, at p1 primordium initiation; right, during further p1 development. Cell clones are numbered (1–12) and outlined (magenta). Leaf primordia (p1–2) are numbered from the youngest. Representative examples from n=23 analysed SAMs are shown. Scale bars, 50 µm.

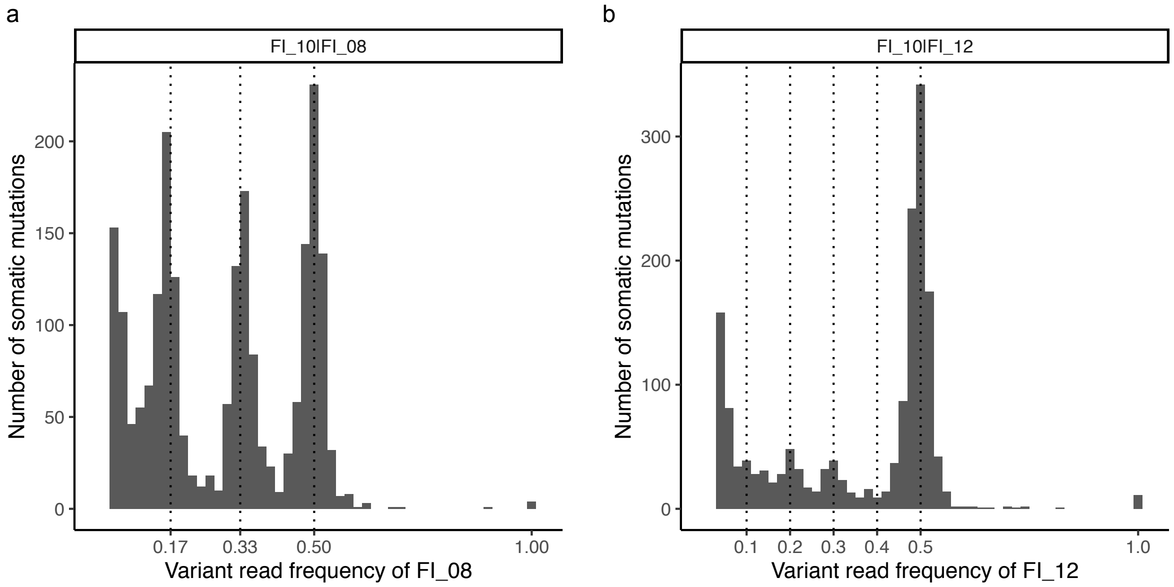

**Supplementary Figure 10: Deep sequencing reveals cell lineages with subfixation of SoGV.** **a,b**, variant read frequency for two ramets with $\sim$ 1370x data from a Finnish clonal lineage at Ängsö. SNPs were called with Mutect2 using a normal-tumor sample pair. For example, "FI_10|FI_0" means FI_10 and FI_08 were used as the normal sample and tumor sample, respectively. Only SNPs meeting the following criteria were kept: 1, Both normal sample and tumor sample have $>$ 100x coverage; 2, Variant read frequency for the normal sample $\geq$ 0.01; 3, Variant read frequency for the tumor sample $\geq$ 0.04. **a**, Histogram of the variant read frequency for FI_08 in the run "FI_10|FI_08". A peak at VRF = 0.5 indicates fixed SoGV with VRF= 0.5. Two additional peaks at 0.17 and 0.33 respectively, indicate sub-fixation. **b**, Histogram of the variant read frequency for FI_12 in the run "FI_10|FI_12". This indicates that there are defined but as yet uncharacterized subpopulations of stem cells where new SoGV are becoming fixed initially, before extending to the entire module.

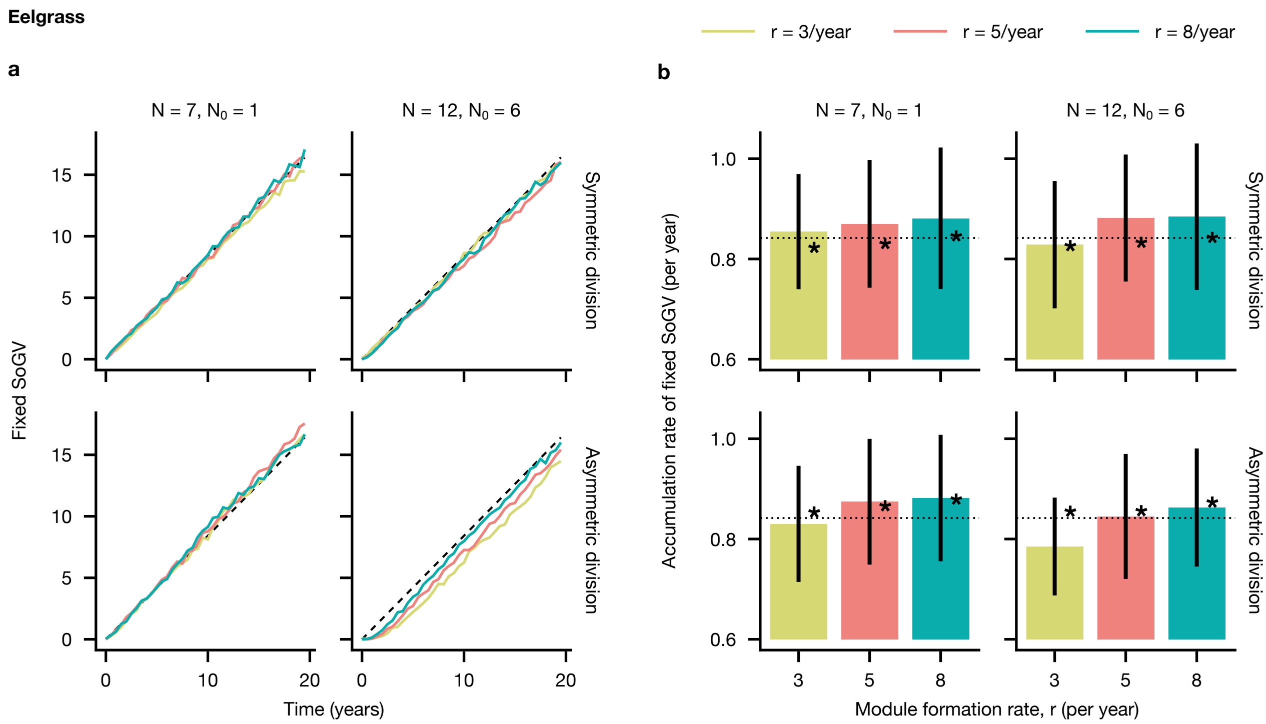

**Supplementary Figure 11:** Agent-based model predictions for the accumulation of fixed somatic mutations in eelgrass via somatic genetic drift with extreme values of $N_0$. The model is parametrized for eelgrass (Supplementary Figs. 7–9). Compared to Figure 3 of the main text, the choice of founder cell size $N_0 = 1$ and $N_0 = 6$ are on the extremes of what is possible for eelgrass. New modules are formed by module branching, which is closer to the biological process in eelgrass. **a**, Data are means over ten simulations. In each simulation the mean fixed mutations are calculated at each time point from a random sample of ten modules. Dashed line: $\mu b$ (approximation of the mutation rate per cell per year). **b**, The accumulation rate of fixed SoGV is estimated from simulated modules at two time points mimicking the experimental methodology (4 years: 3 clones with 2 sampled modules per clone, and 17 years: 2 clones with 5 and 6 sampled modules respectively; see Materials & Methods). Mean fixed SoGV is calculated for each clone and the accumulation rate is then estimated by linear regression. Bars and error bars: mean and standard deviation, respectively, from 100 repeats; dashed line: $\mu b$; stars: accumulation rate of fixed SoGV is estimated by performing linear regression on 100 simulated 200-year old clones and taking the mean. Parameters: $\mu = 0.0069$, $b = 122/yr$, $Z = 1000$ (symmetric update events occur at a rate $b/2$, asymmetric update events at rate $b$).

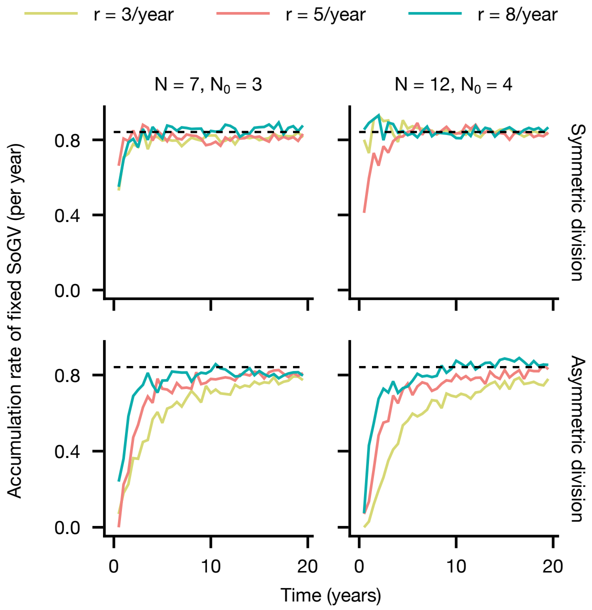

S

**Supplementary Figure 12: Estimated accumulation rates of fixed SoGV calculated from a single time point in simulated eelgrass clones.** Despite different population sizes of (founder) stem cells, i.e. $N$ and $N_0$, accumulation rates are similar. In each simulation the mean number of fixed SoGV is calculated at each time point from a random sample of ten modules and divided by time to obtain the expected rate. Dashed line: $\mu b$ is an approximation of the mutation rate per cell per year. Fixed parameters: $\mu = 0.0069$, $b = 122/yr$, $Z = 1000$ (symmetric update events occur at rate $b/2$, asymmetric update events at rate $b$).

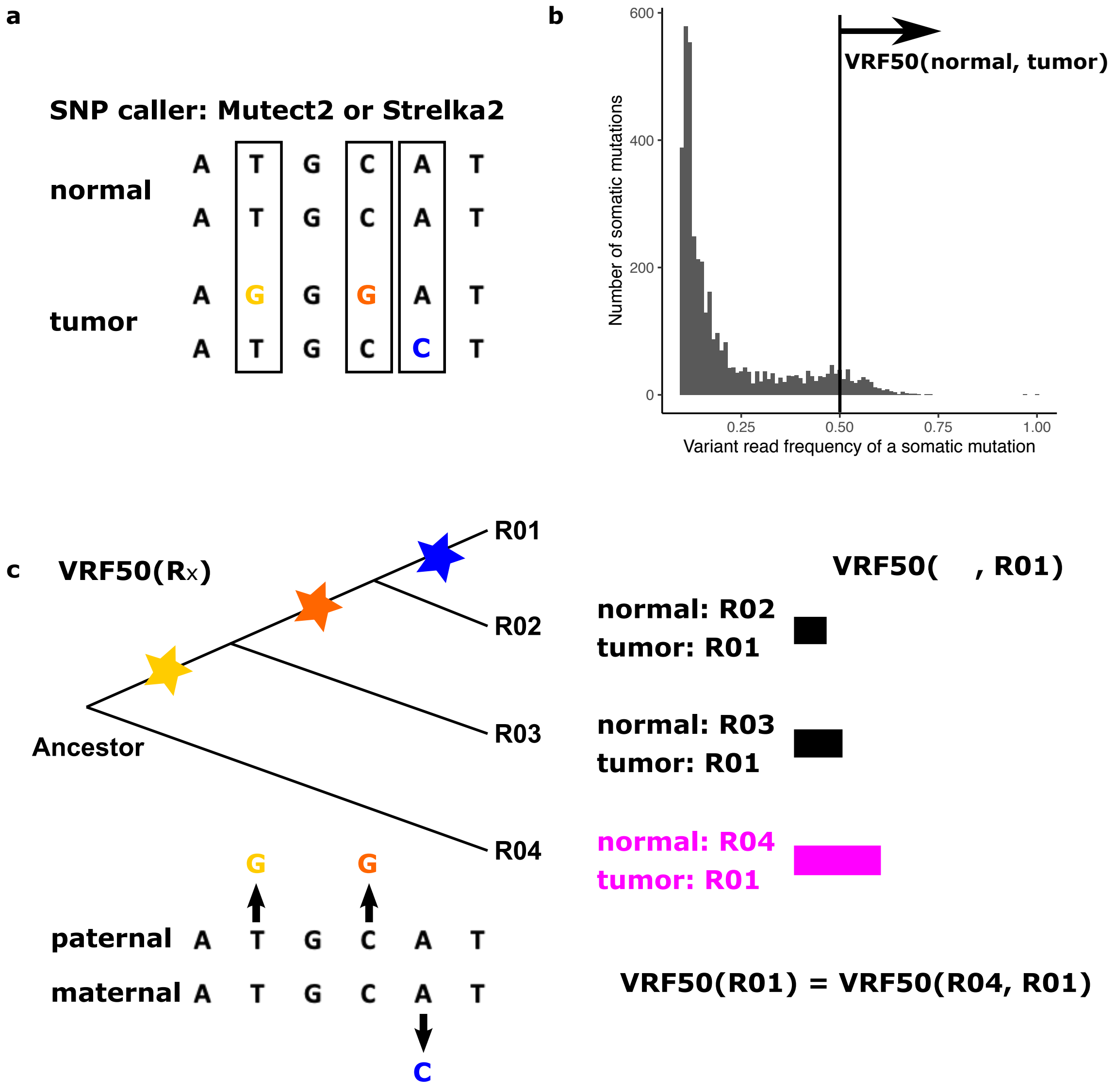

**Supplementary Figure 13: "Variant Read Frequency 50" VRF50($R_x$) as proxy for the number of fixed SoGV. a**, Application of cancer SNP callers. Mutect2 or Strelka2, compares the "normal" sample and the "tumor" sample, and outputs the SNPs where the "normal" sample is homozygous for the reference allele and the "tumor" sample is heterozygous. **b**, VRF50(normal, tumor). For a specific Mutect2/Strelka2 run, VRF50(normal, tumor) represents the number of somatic mutations with a $f_{\mathrm{VRF}} > 0.5$ in the "tumor" sample. **c**, Indirect calculation of VRF50($R_x$). Since the founder seedling/ramet is unavailable, we compare the target ramet $R_x$ with each of the other collected ramets of the same clone, and the maximum value for VRF50(clonemate of $R_x, R_x$) is assigned to VRF50$(R_x)_{\mathrm{obs}}$ (Supplementary Fig. 3). VRF50$(R_x)_{\mathrm{obs}}$ is then standardized based on the proportion of the genome with sufficient coverage to estimate VRF50($R_x$) for the complete genome (Supplementary Data 1).

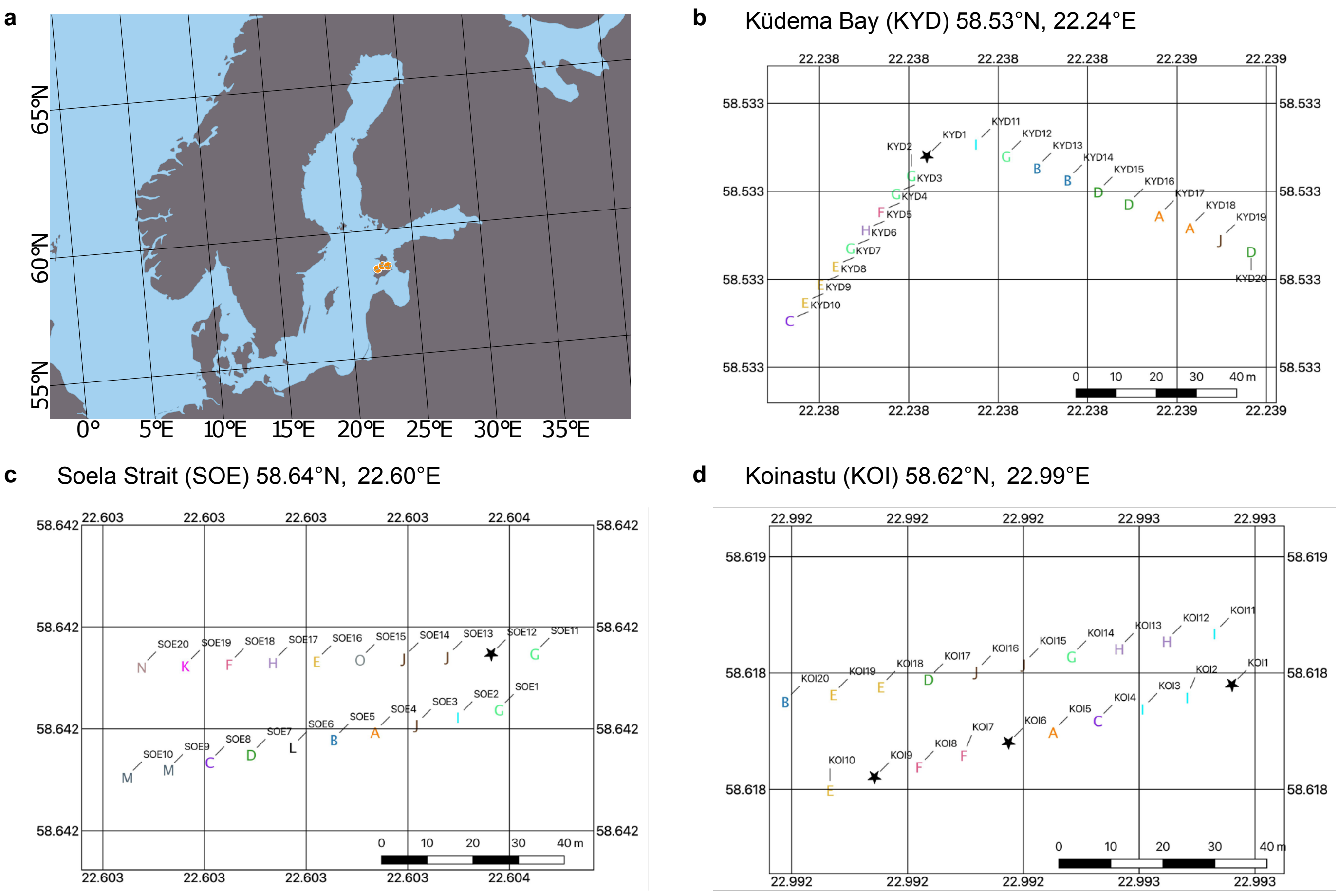

**Supplementary Figure 14: Sampling design for 3 eelgrass *Zostera Marina* populations collected in Estonia, Baltic Sea.** **a**, Overview on sampling sites in the Baltic Sea. **b**, Location Küdema Bay KYD. Each letter indicates a unique clonal lineage, and stars represent missing data. Genets were detected based on 9 microsatellite markers. KYD02, KYD03, KYD06, and KYD12, belonging to genet "G", were used for whole-genome resequencing in this study. **c**, Location Soela Strait SOE. SOE03, SOE13, and SOE14, belonging to clonal lineage "J", were used for whole-genome resequencing in this study. **d**, Location Koinastu KOI. KOI10, KOE18, and KOI19, belonging to genet "E", were used for whole-genome resequencing in this study.
